# Superresolution microscopy reveals partial preassembly and subsequent bending of the clathrin coat during endocytosis

**DOI:** 10.1101/2021.10.12.463947

**Authors:** Markus Mund, Aline Tschanz, Yu-Le Wu, Felix Frey, Johanna L. Mehl, Marko Kaksonen, Ori Avinoam, Ulrich S. Schwarz, Jonas Ries

## Abstract

Eukaryotic cells use clathrin-mediated endocytosis to take up a large range of extracellular cargos. During endocytosis, a clathrin coat forms on the plasma membrane, but it remains controversial when and how it is remodeled into a spherical vesicle.

Here, we use 3D superresolution microscopy to determine the precise geometry of the clathrin coat at large numbers of endocytic sites. Through pseudo-temporal sorting, we determine the average trajectory of clathrin remodeling during endocytosis. We find that clathrin coats assemble first on flat membranes to 50% of the coat area, before they become rapidly and continuously bent, and confirm this mechanism in three cell lines. We introduce the *cooperative curvature model*, which is based on positive feedback for curvature generation. It accurately describes the measured shapes and dynamics of the clathrin coat and could represent a general mechanism for clathrin coat remodeling on the plasma membrane.

## Introduction

Endocytosis is an essential function of eukaryotic cells to internalize molecules from their surface. The major endocytic pathway is clathrin-mediated endocytosis (CME), which supports the uptake of many diverse cargos including nutrients, signaling molecules, membrane proteins, and pathogens. During CME, a dense coat of proteins self-assembles on the inner leaflet of the plasma membrane. The membrane itself is bent into an Ω-shaped invagination that is eventually pinched off to form a coated vesicle, completing endocytosis (Kaksonen and Roux, 2018).

The major component of the coat are clathrin triskelia, which comprise three heavy and three light chains (Fotin et al., 2004). When triskelia are bound to the plasma membrane by adaptor proteins, they form ordered clathrin lattices. The structural flexibility of triskelia allows these lattices to adapt variable ratios of predominantly pentagonal and hexagonal faces, forming both flat and curved geometries.

Both geometries have been observed *in vivo* and *in vitro*, and their structure has been well characterized *in vitro* from a structural biology perspective (Cheng et al., 2007; Dannhauser and Ungewickell, 2012; Fotin et al., 2004; Heuser and Kirchhausen, 1985; Morris et al., 2019; Pearse, 1976; Smith et al., 1998; Takei et al., 1998; Ungewickell and Branton, 1981). However, it remains elusive how clathrin coat formation and membrane bending are temporally and causally related during endocytosis in cells.

In EM micrographs, it was observed early on that clathrin lattices can assume many different curvatures in cells (Heuser, 1980). Since then, two main models of clathrin coat formation during endocytosis have been put forward. In the *constant area model (CAM)*, clathrin grows to its final surface area as a flat coat, which then becomes continuously more curved until vesicle formation is complete. This model assumes that all the differently curved clathrin structures are endocytic intermediates. Early observations suggested the presence of pentagonal and hexagonal faces in isolated coated vesicles (Kanaseki and Kadota, 1969). Combined with the observation of flat lattices being enriched in hexagons, it was suggested that the integration of at least 12 pentagonal faces is a prerequisite for the formation of a spherical structure (Heuser, 1980). However, this would require extensive lattice remodeling which was deemed thermodynamically and structurally unfavorable and thus unlikely to occur (Kirchhausen, 1993). The *constant curvature model (CCM)* was therefore formulated, which assumes that flat clathrin structures are not endocytic precursors. Instead, it was proposed that the endocytic clathrin coat assumes its final curvature, i.e. the curvature of the vesicle, directly from the start, while continuously growing in surface area over time.

The constant curvature model had been prevalent in the field, but was recently challenged by reports that flat clathrin coats can indeed change curvature during endocytosis (Avinoam et al., 2015; Bucher et al., 2018; Scott et al., 2018). This has once again boosted interest in clathrin remodeling (Chen and Schmid, 2020; Kaksonen and Roux, 2018; Sochacki and Taraska, 2018). Recent years have seen numerous studies based on diverse methods that did not converge on a common result. Reports supported either the constant area model (Avinoam et al., 2015; Sochacki et al., 2021), the constant curvature model (Willy et al., 2021), a combination of both models (Bucher et al., 2018; Tagiltsev et al., 2021; Yoshida et al., 2018) or the simultaneous existence of both models within cells (Scott et al., 2018). To conclusively understand this complex process, it would be desirable to directly visualize the three-dimensional nanoscale structure of the endocytic clathrin coat in living cells. Unfortunately, however, to date no method offers the necessary spatial and temporal resolution to do that. Here, we aim to circumvent this limitation by densely sampling the entire endocytic process in fixed cells, and subsequently reconstructing the dynamic information.

We developed a superresolution microscopy approach to quantitatively study the three-dimensional (3D) clathrin coat architecture at endocytic sites. We used a novel model fitting framework to extract geometric parameters that allowed us to sort static images of clathrin lattices according to their progression along the endocytic timeline. The inferred endocytic dynamics allowed us to reconstruct the stereotypic remodeling of clathrin during endocytosis at the nanoscale.

In summary, we found that a clathrin coat first partially assembles on a flat membrane, and then simultaneously grows in surface area and coat curvature. While initial bending occurs rapidly, it later slows down and endocytic sites are eventually paused in a state of high curvature before vesicle scission. This trend is conserved across cell lines, suggesting it is a common attribute that is not affected by cell type. Based on this data, we developed a new kinetic growth model, the *cooperative curvature model* (*CoopCM*). It describes coat area growth as addition of clathrin to the lattice edge with a constant rate and assumes that the curvature of the coat increases towards a preferred curvature, driven by the cooperative interplay of clathrin triskelia within the lattice. The CoopCM predicts a fast initial curvature increase that is slowed down progressively as the coat becomes spherical, and shows excellent agreement with experimental data.

## Results

### Quantitative 3D superresolution imaging of clathrin structures

Here, we used 3D single-molecule localization microscopy (SMLM) to systematically determine the precise geometry of individual clathrin-coated structures at the plasma membrane. For this, we optimized the sample preparation to label clathrin at endocytic sites as densely as possible using indirect immunofluorescence with polyclonal antibodies against clathrin light and heavy chains, which was crucial to achieve high quality SMLM (Mund and Ries, 2020). We then localized sparsely activated single fluorophores by fitting an experimentally derived model of the astigmatic point-spread function using a method we developed recently (Li et al., 2018). This improved the resolution to about 10 nm in x/y and 30 nm in z (based on modal values of the localization precision at 3.9 nm in x/y and 12.5 nm in z, see Methods), and reduced typically observed image distortions. The achieved image quality allowed us to clearly visualize and quantify the 3D clathrin coat shape at different stages of endocytic site maturation (Figure 1A-C).

**Figure 1.**
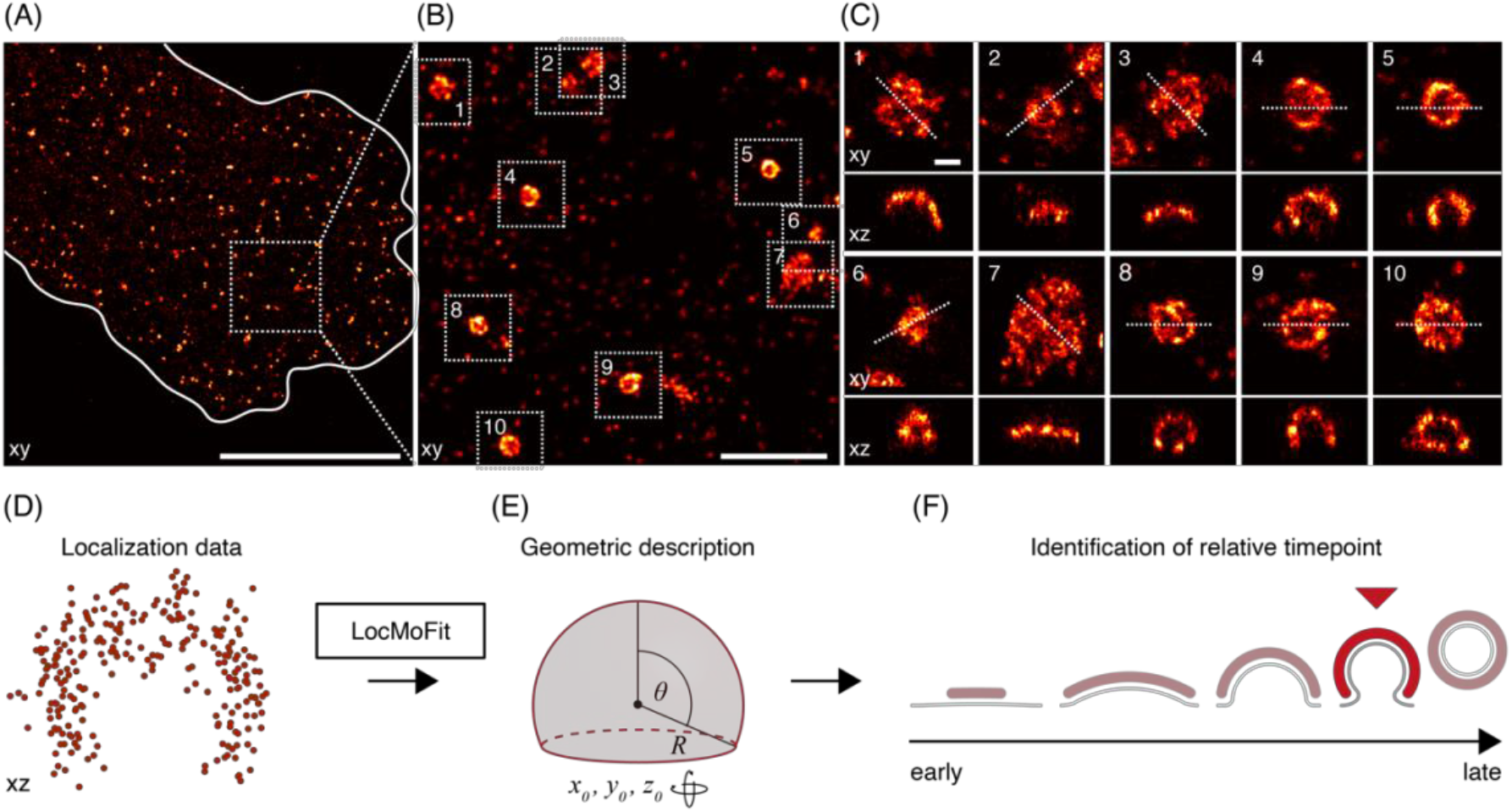
3D SMLM of clathrin-coated structures. (A) 3D SMLM image of clathrin immunolabeled with AF647 at the bottom membrane of a SK-MEL-2 cell. (B) Enlarged view of the region indicated in (A). (C) All structures found in (B) are shown in top view (xy) and as 50 nm-thick z-slices (xz), orientation of slice indicated by dotted line). Scale bars are 10 μm (A); 1 μm (B); and 100 nm (C). (D) Geometric analysis pipeline. All clathrin coats are segmented, and their localizations are directly fitted with a spherical cap model using the fitting framework LocMoFit (Wu et al., 2021). (E) In LocMoFit, individual clathrin coats are parametrized by their size (radius R), closing angle (θ), position (x_0_, y_0_, z_0_) and 3D orientation. (E) Using θ as a proxy for endocytic progression, the relative endocytic time point for each structure can be determined.

The large majority of sites were single structures that were well-isolated from each other, and exhibited great structural diversity. In addition, we noted several clusters of closely juxtaposed endocytic sites (Supplementary Figure 1A). In the isolated sites, we observe a variety of shapes including flat, curved, dome-shaped and spherical structures of different sizes (Figure 1C), indicating that endocytosis has been arrested by fixation at different time points during endocytic progression.

To quantify size and shape of individual clathrin coats, we used LocMoFit, a computational pipeline based on a maximum-likelihood model fitting framework that we developed previously (Wu et al., 2021). This framework directly fits 3D geometric models to the 3D point cloud of localizations (Figure 1D-E) instead of rendered images, and thereby harnesses the full information contained in the SMLM data, including for instance the localization precision, rather than just spatial coordinates.

We describe the clathrin coat as a spherical cap that is defined by a radius *r* and a closing angle *θ* (Figure 1E). Our model also accounted for coat thickness, antibody size and blurring due to the localization precision (Methods). Hence, it describes both shallow and highly curved structures equally well. Moreover, because *θ* increases during endocytosis from 0° at flat sites to 180° at complete vesicles, we could use this parameter to sort the individual images according to endocytic progress (Figure 1D).

### The clathrin coat grows in area and becomes more curved during endocytosis

We first imaged immunostained clathrin in chemically fixed SK-MEL-2 cells, where we focused on the bottom plasma membrane that was densely covered by endocytic sites. These cells have been extensively studied and are established model cells for clathrin-mediated endocytosis with well-characterized endocytic dynamics (Aguet et al., 2013; Avinoam et al., 2015; Doyon et al., 2011; Kaplan et al., 2020). Using the 3D model fitting pipeline, we determined radius *R*, closing angle *θ*, position and rotation of 1798 endocytic sites from 13 cells (Figure 2A) with high accuracy (Supplementary Figure 2). We found that two structural intermediates are enriched, while others are rare and only a small fraction of sites were completely flat (Figure 2B). Slightly curved sites with *θ* ≈ 70° and strongly curved sites with *θ* ≈ 130° were enriched, indicating that those structural states are more frequently found in cells. We only rarely obtained sites with *θ* ≈ 180°, which would be expected for complete spherical coats, even though fully formed vesicles are found in our data (Supplementary Figure 1B). This indicates that at the time point of scission, the clathrin coat of nascent vesicles is still incomplete at the neck, or that the kinetics of scission are too transient for detection. Deeply invaginated clathrin coated pits could further be sheared off during sample preparation, thus evading our detection. Curvatures *H* = 1/*R* ranged from 0 to 0.022 nm^-1^ with a median of 0.011 nm^-1^, corresponding to a median radius of 87 nm, and surface areas ranged from 9,000 nm^2^ to 140,000 nm^2^ with a median of 54,000 nm^2^. These values agree well with previous measurements of the vesicular coat using EM (Avinoam et al., 2015), median curvature 0.015 nm^-1^, median surface 54,500 nm^2^).

**Figure 2.**
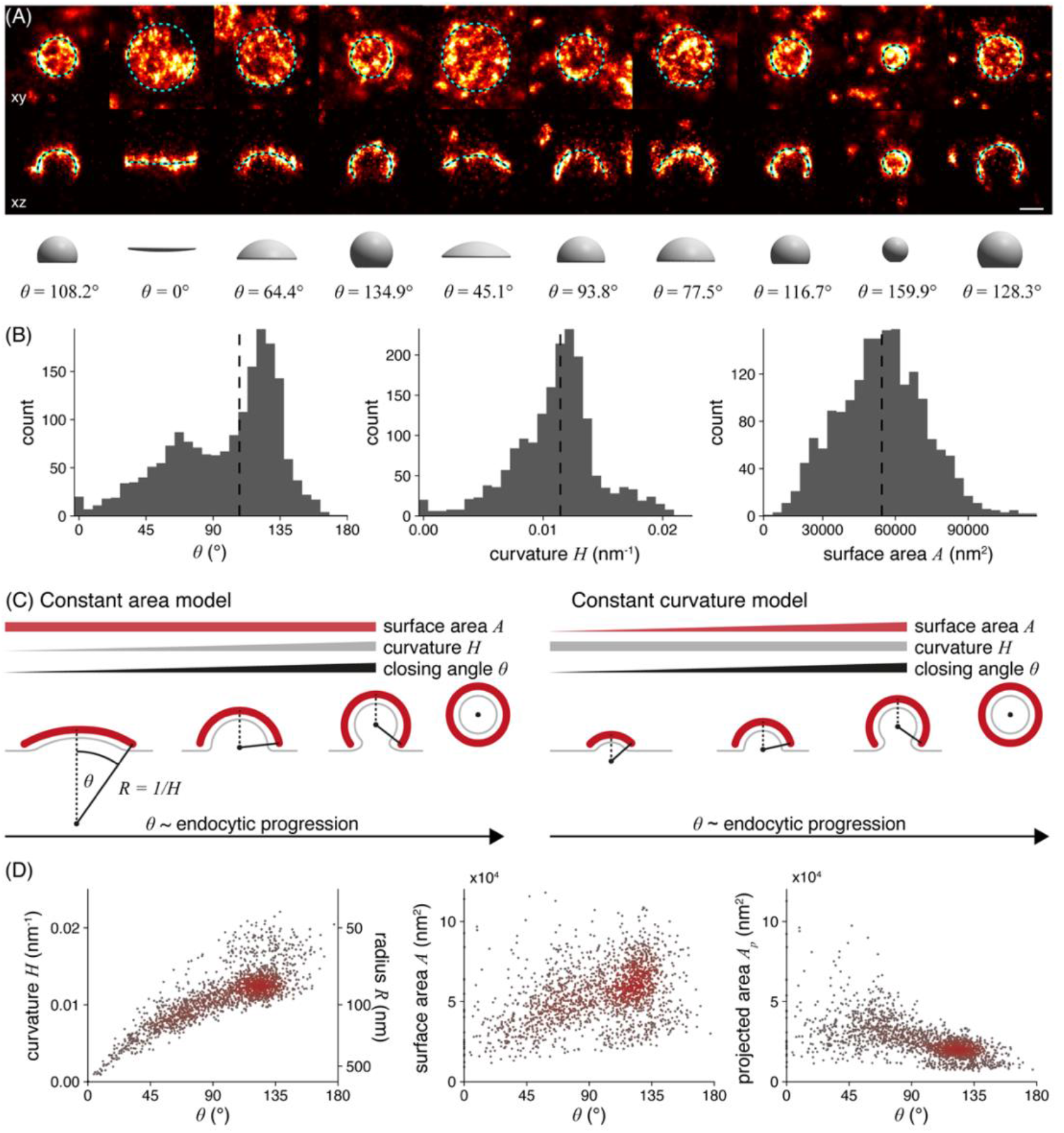
Quantitative analysis of clathrin-coated structures in SK-MEL-2-cells. (A) Clathrin coat geometry is quantified using LocMoFit. The fitted spherical cap model is drawn as circle with its outer radius (top row, xy), as cross-sectional profile (50 nm xz slices, middle row), and as surface with the corresponding closing angle θ (bottom row). Scale bar 100 nm. (B) Distributions of closing angle θ (median = 108°), curvature H (median = 0.011 nm^-1^) and surface area A (median = 54,000 nm^2^) of endocytic sites in SK-MEL-2 cells (n = 1798 sites, N = 13 cells) as determined from the model fit. (C) Two previously proposed mechanistic models for clathrin coat assembly during endocytosis (for details see text). In both scenarios θ increases monotonically and is thus a proxy for endocytic progression. (D) Development of curvature, surface area and projected area of the clathrin coat during endocytosis. Color indicates point density.

We then wanted to understand how the clathrin coat geometry changes during endocytosis. To this end, we used the closing angle parameter *θ* to sort endocytic sites relative to each other in time. Irrespective of whether endocytosis follows a constant curvature or constant area model, *θ* monotonically increases from early to late endocytic time points (Figure 2C), and can thus be used as unbiased proxy for endocytic progression (Avinoam et al., 2015).

The curvature *H* was strongly correlated with *θ*, indicating that coat curvature increases continuously during endocytosis (Figure 2D). Similarly, the surface area increased from 32,000 nm^2^ (median of 5% of sites with smallest theta) to 50,000 nm^2^ (median of 5% of sites with highest theta). The projected area decreased from 31,000 nm^2^ to 13,000 nm^2^ (median of 5% of sites with lowest and highest theta respectively), which is in close agreement with previous EM measurements (Bucher et al., 2018). It is readily obvious that our data are incompatible with the constant curvature model, as the curvature is not constant, but increases monotonically with *θ*. Just as clearly, our data do not support the constant area model, because the coat surface also increases during endocytosis.

Almost all data points are part of one continuous point cloud (Figure 2D), indicating a continuous transition between structural states during endocytosis. We noticed an additional small, disconnected set of data points representing 8.5% of all sites that correspond to endocytic sites with curvatures above 0.016 nm^-1^ and *θ* of 80°-180° (Figure 2D, and example structures in Supplementary Figure 3). In a control experiment to check whether these are endocytic structures, we selectively analyzed clathrin structures that colocalized with AP-2, a bona fide marker for CME. We did not observe AP-2 in any of the disconnected sites, from which we conclude that they indeed do not belong to CME. These small structures could represent coated vesicles from the *trans* Golgi (Supplementary Figure 4).

### The cooperative curvature model of clathrin coat remodeling

Our data clearly showed that clathrin coats grow and become more curved as the plasma membrane gets bent to produce a vesicle. Because the data quantitatively describes all 3D geometries that the endocytic clathrin coat assumes, it allowed us to move towards a mathematical model of clathrin coat formation during endocytosis.

Here, we introduce the *Cooperative Curvature Model* (*CoopCM*), which describes coat growth based on known structural and dynamical properties of clathrin coats (Figure 3A). Firstly, we assume that the clathrin coat starts growing on a flat membrane. While triskelia have been shown to exchange dynamically during endocytosis (Avinoam et al., 2015), net growth of the area *A* occurs via the addition of triskelia at the lattice edge *Ɛ* with a constant growth rate *k*_on_ (Equation 1).

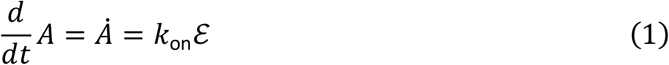

**Figure 3.**
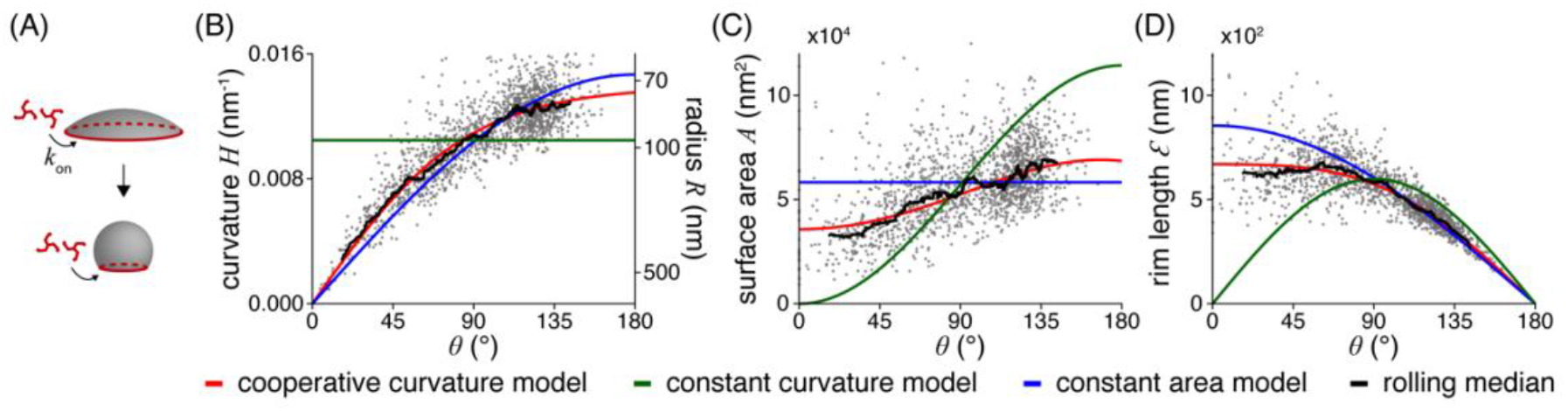
Model for clathrin coat growth. (A) Schematic of the cooperative curvature model, where clathrin lattices grow by addition of triskelia to the rim at a constant growth rate k_on_. Curvature generation is driven towards a preferred curvature, ultimately creating a spherical vesicle. (B) Distinct clathrin growth models and rolling median (window width = 82 sites) fitted to curvature over θ. The resulting fitting parameters are then used to map the same models also over (C) surface area and (D) rim length. (n = 1645 sites, N = 13 cells).

Moreover, we assume that the intrinsic pucker angle of individual triskelia and their interactions in a lattice together translate into an overall preferred curvature *H*_0_ of the clathrin coat as a whole. However, the initially flat assembly suggests that this preferred curvature cannot be realized in an immature lattice. We hypothesize that cooperative processes in the coat are required to increase curvature. Coat curvature *H* then increases asymptotically towards *H*_0_ at an initial rate *γ*, slowing down towards zero when the preferred curvature is reached (Equation 2).

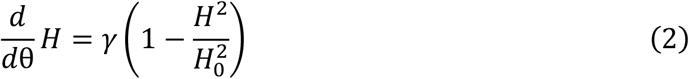

Choosing a quadratic dependence of the rate of curvature change on curvature is a simple way to represent cooperativity in the lattice, which recently has been demonstrated in experiments (Sochacki et al., 2021; Zeno et al., 2021) and fits our data more accurately than the linear relationship, which would correspond to a less cooperative process (Supplementary Note). Equation 2 can be solved analytically to yield an expression of the curvature *H* depending on *θ* (Equation 3), which can then be fitted to our data.

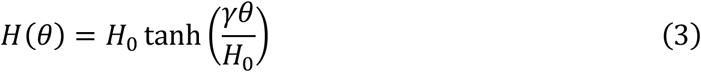

Analogous expressions for other geometric parameters like surface area can be derived straightforwardly.

The fit of this CoopCM to the data of curvature *H* in relation to closing angle *θ* shows excellent agreement (Figure 3B). In comparison, the CAM fitted the curvature data slightly worse than the CoopCM, and the constant curvature model agreed poorly with the data. The improved fit of the CoomCM compared to the CAM or CCM is not a result of the additional fitting parameter, as we showed using the Baysian Information Criterion (see Supplementary Note).

From the curvature fits, we calculated the corresponding curves for surface area (Figure 3C) and the edge length of the clathrin coat (Figure 3D). The edge length decreases monotonically and approaches zero at *θ* = 180°, thereby stopping growth according to Equation 1. Both graphs again highlight the very close agreement of the model prediction with the experimental data. From the fit, we determined that invagination occurs when about half of the total coat area has grown (*A*_0_ = 0.51), and that the preferred curvature of a clathrin coat is *H*_0_ = 0.014 nm^-1^, corresponding to a radius of *R*_0_ = 72 nm. The model yields nearly identical parameters when surface area or rim length are fitted instead of curvature (Table 1), highlighting its robustness.

**Table 1.** Summary of clathrin coat growth model fits in SK-MEL-2. Fitted parameter values for the constant area model (CAM), the constant curvature model (CCM) and the cooperative curvature model (CoopCM) when fitting curvature H(θ), surface area A(θ), rim length Ɛ(θ) and projected area A_p_(θ). A: Surface area fitted with CAM; R: Radius fitted with CCM; γ: Constant rate of curvature increase fitted with CoopCM; H_0_: Preferred curvature of the clathrin coat fitted with CoopCM; R0: preferred radius of the clathrin coat fitted with CoopCM; A_0_: Fraction of surface area growing as a flat lattice before curvature initiation, defined as A_0_ = A(θ = 0.01)/A(θ = π); k_on_: local growth rate obtained from θ (t) fitted with CoopCM and measured in nm per pseudotime units 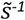. N = 13 cells; n = 1645 sites.

| | | Curvature $H(\theta)$ | Surface area $A(\theta)$ | Rim length $\mathcal{E}(\theta)$ | Projected area $A_p(\theta)$ |
| --- | --- | --- | --- | --- | --- |
| **CAM** | **A** | $58300 \pm 400 \text{ nm}^2$ | $56600 \pm 400 \text{ nm}^2$ | $49400 \pm 500 \text{ nm}^2$ | $48800 \pm 500 \text{ nm}^2$ |
| **CCM** | **R** | $94.5 \pm 0.7 \text{ nm}$ | $84.8 \pm 0.4 \text{ nm}$ | $97.3 \pm 0.8 \text{ nm}$ | $92.8 \pm 0.7 \text{ nm}$ |
| **CoopCM** | **$\gamma$** | $(9.4 \pm 0.1) \cdot 10^{-3} \text{ nm}^{-1}$ | $(9.0 \pm 0.1) \cdot 10^{-3} \text{ nm}^{-1}$ | $(9.4 \pm 0.1) \cdot 10^{-3} \text{ nm}^{-1}$ | $(9.1 \pm 0.1) \cdot 10^{-3} \text{ nm}^{-1}$ |
| | **$H_0$** | $(13.9 \pm 0.1) \cdot 10^{-3} \text{ nm}^{-1}$ | $(13.8 \pm 0.1) \cdot 10^{-3} \text{ nm}^{-1}$ | $(13.4 \pm 0.1) \cdot 10^{-3} \text{ nm}^{-1}$ | $(13.6 \pm 0.1) \cdot 10^{-3} \text{ nm}^{-1}$ |
| | **$R_0$** | $72.0 \pm 0.6 \text{ nm}$ | $72.2 \pm 0.6 \text{ nm}$ | $74 \pm 0.8 \text{ nm}$ | $73.4 \pm 0.9 \text{ nm}$ |
|  | **$A_0$** | 0.51 | 0.55 | 0.49 | 0.52 |
| | **$k_{on}$** | $78.1 \text{ nm}\tilde{s}^{-1}$ | $72.2 \text{ nm}\tilde{s}^{-1}$ | $83.4 \text{ nm}\tilde{s}^{-1}$ | $78 \text{ nm}\tilde{s}^{-1}$ |

We then decided to test if the observed mode of clathrin remodeling is specific for SK-MEL-2 cells. For this, we analyzed the endocytic clathrin ultrastructure also in U2OS cells and 3T3 mouse fibroblasts (Supplementary Figure 5 and Table 2-3). In these cell lines, just like in SK-MEL-2 cells, the curvature as well as the surface area continuously increase throughout endocytosis. We observe that the preferred curvature of the clathrin coat is smaller in U2OS (*H*_0_ = 0.012 nm^-1^, *R*_0_ = 85 nm) and 3T3 cells (*H*_0_ = 0.011 nm^-1^, *R*_0_ = 89.7 nm) compared to SK-MEL-2 (*H*_0_ = 0.014 nm^-1^, *R*_0_ = 72 nm). This suggests that the average size of vesicles formed in CME is cell line specific. The fraction of the surface area acquired on the flat membrane is very similar for all three cell lines, with U2OS derived sites initiating curvature at *A*_0_ = 0.52 and 3T3 sites at *A*_0_ = 0.45 (Table 2-3). Taken together, we have shown in several cell lines that clathrin coats neither grow via constant curvature or constant area pathways, but rather first grow flat and then acquire curvature and more surface area simultaneously, with a non-linear mode of curvature generation.

**Table 2.** Summary of clathrin coat growth model fits in U2OS. Fitted parameter values for the constant area model (CAM), the constant curvature model (CCM) and the cooperative curvature model (CoopCM) when fitting curvature H(θ), surface area A(θ), rim length Ɛ(θ) and projected area A_p_ (θ). A: Surface area fitted with CAM; R: Radius fitted with CCM; γ: Constant rate of curvature increase fitted with CoopCM; H_0_: Preferred curvature of the clathrin coat fitted with CoopCM; R0: preferred radius of the clathrin coat fitted with CoopCM; A_0_: Fraction of surface area growing as a flat lattice before curvature initiation, defined as A_0_ = A(θ = 0.01)/A(θ = π); *k*_on_: local growth rate obtained from θ (t) fitted with CoopCM and measured in nm per pseudotime units 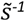. N = 3 cells; n = 241 sites.

| | | Curvature $H(\theta)$ | Surface area $A(\theta)$ | Rim length $\mathcal{E}(\theta)$ | Projected area $A_p(\theta)$ |
| --- | --- | --- | --- | --- | --- |
| **CAM** | **A** | $80800 \pm 1600 \text{ nm}^2$ | $76900 \pm 1700 \text{ nm}^2$ | $65900 \pm 1600 \text{ nm}^2$ | $65400 \pm 1600 \text{ nm}^2$ |
| **CCM** | **R** | $119.1 \pm 2.5 \text{ nm}$ | $101.6 \pm 1.5 \text{ nm}$ | $122.8 \pm 2.7 \text{ nm}$ | $115 \pm 2.4 \text{ nm}$ |
| **CoopCM** | **$\gamma$** | $(7.9 \pm 0.2) \cdot 10^{-3} \text{ nm}^{-1}$ | $(7.7 \pm 0.2) \cdot 10^{-3} \text{ nm}^{-1}$ | $(8.1 \pm 0.2) \cdot 10^{-3} \text{ nm}^{-1}$ | $(7.9 \pm 0.2) \cdot 10^{-3} \text{ nm}^{-1}$ |
| | **$H_0$** | $(11.8 \pm 0.2) \cdot 10^{-3} \text{ nm}^{-1}$ | $(11.5 \pm 0.2) \cdot 10^{-3} \text{ nm}^{-1}$ | $(11.0 \pm 0.3) \cdot 10^{-3} \text{ nm}^{-1}$ | $(11.1 \pm 0.3) \cdot 10^{-3} \text{ nm}^{-1}$ |
| | **$R_0$** | $85.0 \pm 1.8 \text{ nm}$ | $87.1 \pm 1.7 \text{ nm}$ | $90.5 \pm 2.7 \text{ nm}$ | $90.2 \pm 2.6 \text{ nm}$ |
|  | **$A_0$** | 0.52 | 0.52 | 0.45 | 0.47 |
| | **$k_{on}$** | $89.3 \text{ nm}\tilde{s}^{-1}$ | $91.5 \text{ nm}\tilde{s}^{-1}$ | $110.3 \text{ nm}\tilde{s}^{-1}$ | $105.5 \text{ nm}\tilde{s}^{-1}$ |

**Table 3.**
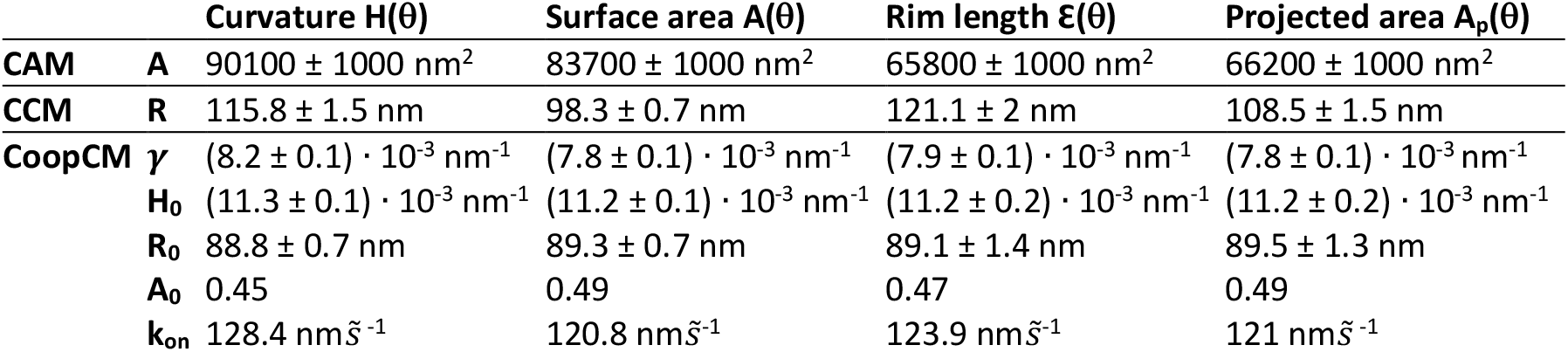
Summary of clathrin coat growth model fits in 3T3 mouse fibroblasts. Fitted parameter values for the constant area model (CAM), the constant curvature model (CCM) and the cooperative curvature model (CoopCM) when fitting curvature H(θ), surface area A(θ), rim length Ɛ(θ) and projected area A_p_(θ). A: Surface area fitted with CAM; R: Radius fitted with CCM; γ: Constant rate of curvature increase fitted with CoopCM; H_0_: Preferred curvature of the clathrin coat fitted with CoopCM; R_0_: preferred radius of the clathrin coat fitted with CoopCM; A_0_: Fraction of surface area growing as a flat lattice before curvature initiation, defined as A_0_ = A(θ = 0.01)/A(θ = π); k_on_: local growth rate obtained from θ (t) fitted with CoopCM and measured in nm per pseudotime units 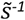. N = 7 cells; n = 688 sites.

### Temporal reconstruction of structural dynamics in endocytosis

We systematically segment all endocytic sites in the superresolution images, and thereby obtain a large dataset where all endocytic time points are sampled homogenously. The distribution of structural states within the dataset is thus representative of their lifetime, with long-lived, stable curvature states being overrepresented in the data compared to transient states. This opens up the possibility to reconstruct the temporal progression of clathrin remodeling during endocytosis, and to ask whether clathrin coats acquire their curvatures with a constant rate, or if there are certain curvature transitions that occur more rapidly than others.

For this, we sorted all endocytic sites by *θ*. The rank of an endocytic site thus corresponds to its pseudotime, which describes its relative time point between 0 and 1 along the endocytic trajectory (Figure 4A).

**Figure 4.**
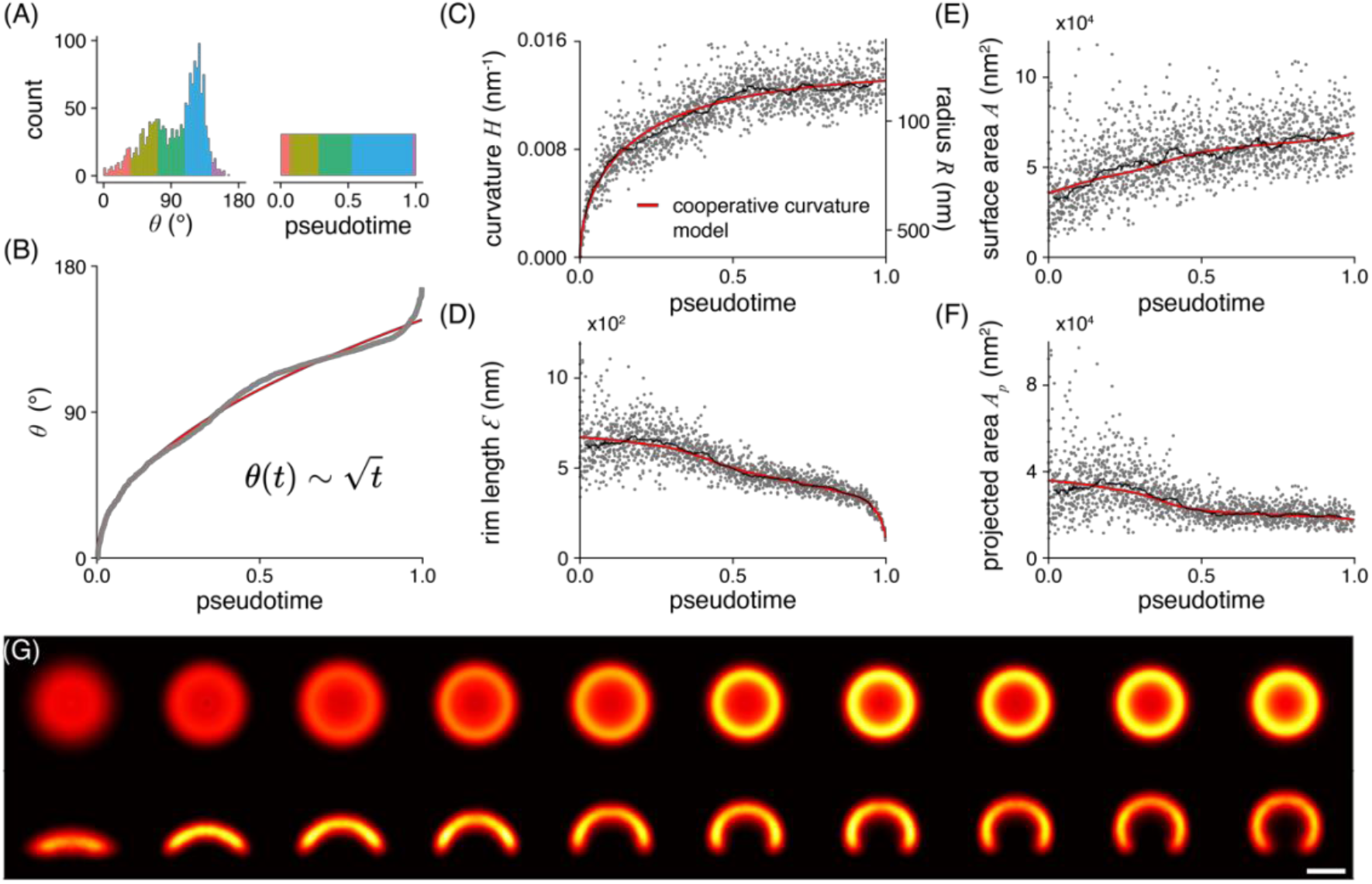
Temporal reconstruction of clathrin coat remodeling. (A) Endocytic sites are sorted by θ to reconstruct pseudotime. Enriched θ states, for example the peak at 135°, represent long-lived states that remain for extended periods in pseudotime. Color code represents the same number of sites in both histograms. (B) The square-root dependence between θ and pseudotime approximated by the cooperative curvature model (red line). (C) Curvature generation over pseudotime is approximated by the cooperative curvature model. Fit results in (C) were used to describe (D) rim length, (E) surface area and (F) projected area change over pseudotime. A rolling median (window of 82 sites) is plotted alongside (black line). (G) Superresolution averages for distinct endocytic stages, resulting from all collected snapshots. Each bin contains the same number of snapshots of clathrin-coated structures sorted along their pseudotime (n = 163 per bin), so that all bins represent equally long pseudotime intervals. Individual sites were rescaled to the median bin radius and aligned by their center coordinates as well as rotation angles. Scale bar is 100 nm. (n = 1645 sites, N = 13 cells).

As our model (Equation 1) describes the dynamic growth of the clathrin coat, we can solve it to derive an expression for *θ* over time *t* (Equation 4), where the coat starts out flat and then starts to generate curvature, increasing *θ* over time.

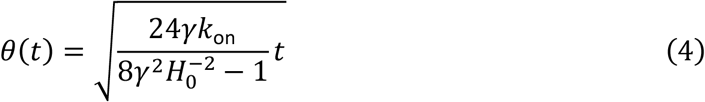

The square root dependence of *θ* on time t reflects the slowing down of curvature generation as the clathrin coat approaches its preferred curvature. This expression fits the pseudotime-resolved data remarkably well (Figure 4B). Consistent with our previous reasoning, a linear model did not agree well with the data (Supplementary Note), since it gives a linear invagination speed for small *t*, emphasizing the validity of our cooperative curvature model, which leads to the characteristic square root dependence. The data slightly deviates from the model in the early phase, and more notably in the very end for pseudotimes close to 1. This potentially indicates that clathrin geometry is influenced by other factors besides the coat itself close to vesicle scission.

Because this pseudotime resolved dataset was generated from a large number of endocytic sites in many cells, we effectively generated the average trajectories of how curvature, surface area, projected area and lattice edge change during endocytosis in SK-MEL-2 cells (Figure 4C-F). We observe comparatively few flat clathrin coats. This shows that clathrin lattices are only transiently flat in the beginning of endocytosis, and might represent an energetically unfavorable conformation (Figure 4B). A fast transitions from a flat to a curved structure could further be mediated independently of clathrin coat (Zhao et al., 2017), e.g. by the action of BAR domain proteins (Henne et al., 2010), or at the base of filopodial projections. We further find comparatively many structures with a curvature of *θ* ≈ 70° and *θ* ≈ 130°, indicating more long-lived endocytic stages. As the surface area is constantly increasing over pseudotime, we assume that the enrichment at *θ* ≈ 130° represents a stalling point, where vesicles are formed by addition of final clathrin triskelia and potentially other factors that enable the recruitment and mechanic function of dynamin during vesicle scission. Similarly, the enrichment at 0 ≈ 70° could be indicative of further recruitment of regulatory components, potentially supporting a previously suggested endocytic checkpoint (Loerke et al., 2009). This functional interpretation is supported by the observation of similar peaks in U2OS (at *θ* ≈ 60° and ≈ 130°) as well as 3T3 cells (at *θ ≈* 50° and ≈ 130°).

Within the first 10% of pseudotemporal progression, the clathrin coat rapidly acquires shallow curvature of *H* = 0.007 nm^-1^ (*R* = 134 nm). It then becomes gradually more bent up to ≈60% of the endocytic timeline, when the sites reach an average curvature of 0.012 nm^-1^ (*R* = 83 nm). During the last ≈40% of progression, the curvature increase almost stops, until vesicle scission occurs. Interestingly, the earliest sites in our dataset already contain ≈50 % of the surface area of the final vesicles, which is also reflected in the fitting results (*A*_0_ = 0.51) (Figure 4E). This indicates that the initial assembly of the clathrin coat occurs very rapidly, or is in part below our detection limit.

We performed identical analyses for U2OS and 3T3 derived clathrin sites and obtained highly similar results as for SK-MEL-2 cells, indicating that this trajectory represents a general pathway for clathrin coat remodeling during endocytosis (Supplementary Figure 5 and Tables 2-3). Finally, we made a 3D nanoscale movie, which vividly illustrates assembly and remodeling of the clathrin coat during endocytosis, from pseudotime resolved averages of our data (Figure 4G). This yielded a nanoscale pseudo-temporal movie of endocytosis in SK-MEL-2 cells (Supplementary Movie).

## Discussion

### Quantitative description of clathrin coat ultrastructure

The nature of clathrin assembly during endocytosis has become a classical question in cell biology that remains unresolved (reviewed in (Chen and Schmid, 2020; Sochacki and Taraska, 2018)). There have been two main competing mechanistic models of how triskelia form a coat: the constant curvature model predicts that the clathrin coat assumes its final curvature directly from the start, while continuously increasing in surface area over time. In contrast, the constant area model predicts that clathrin grows to its final surface area as a flat coat, which then becomes continuously more curved until vesicle formation is complete.

Each model is supported by a number of studies, which mostly relied on electron microscopy techniques. Among them, 3D correlative electron tomography yields exquisite 3D endocytic membrane shapes (Avinoam et al., 2015), but can be tedious to extend towards large numbers of cells. Platinum replica electron microscopy offers large fields of view and thus superb throughput, but gives limited 3D resolution (Bucher et al., 2018; Sochacki et al., 2017).

We reason that the trajectory of clathrin coat assembly during endocytosis could best be determined by systematically imaging large numbers of endocytic sites with high 3D resolution. Therefore, we used SMLM that combines high 3D resolution with high throughput. Although lower than in electron microscopy, the resolution we achieved allowed us to resolve the precise 3D coat ultrastructure of clathrin-coated pits. Importantly, due to the molecular specificity for clathrin, we were able to segment all endocytic sites at the bottom surface of the cells in our images in an unbiased way, thus ensuring homogenous sampling of the structural variety of clathrin coats.

We applied a novel maximum-likelihood based fitting framework that we developed recently (Wu et al., 2021), which allows fitting complex geometric models to localization point clouds. Because the fit is applied directly to the localizations, it considers all quantitative parameters of all localizations, most notably localization uncertainty. This results in the precise and reliable quantification of the underlying structures (Wu, Tschanz & Krupnik et al., 2020), even when taking linkage error through indirect immunolabeling into account (Supplementary Figure 6).

We described the clathrin coat as a spherical cap, which matched the large majority of sites very well. Additionally, in our data, we observed that some sites are asymmetric, ellipsoidal, or deformed more irregularly (Supplementary Figure 1C). We do not currently evaluate this asymmetry. Rather, we introduced rotational symmetry during the averaging (Figure 4G), where we aligned each site based on their model parameters and thus averaged out many irregularities. A deviation of the cross-sectional profile from a circle is nevertheless preserved in the averaging (Supplementary Figure 8D - F), and in future studies more complex geometric models will enable analyzing the structure and asymmetries of the coat in more detail.

### Flat clathrin-coated structures

In the first step of our analysis pipeline, we segmented all clathrin-coated structures in the superresolution images. We thereby aimed to achieve homogenous sampling to capture all endocytic intermediates with the same probability irrespective of time point, size, and orientation. Interestingly, the surfaces of the earliest detectable endocytic sites already contained half the surface area of sites in the latest CME stages. How do these earliest sites assemble? We cannot sort the very flat sites, which are comparatively rare in our dataset, in time using *θ* because their *θ* is close to zero. However, their relative fraction among all sites is still informative. As short-lived states are under-represented, the relative absence of small flat sites in our data indicates that the initial assembly occurs rapidly.

In addition to the short lifetimes of small flat sites, two technical reasons could potentially contribute to their rare occurrence in our data. Due to their small size and potentially sub-stoichiometric labeling, which is also noticeable as holes in larger structures (Supplementary Figure 1D), they might not have sufficient immunostaining signal and thus they might be hard to be differentiated from small clusters of primary and secondary antibodies that are routinely observed in SMLM images. Additionally, the very first small clathrin assemblies might be meta-stable, with mostly loosely bound triskelia, and thus might not be stabilized well by chemical fixation. However, it is important to note that very small sites are also rarely found in previously published electron microscopy datasets (Avinoam et al., 2015; Bucher et al., 2018; Sochacki et al., 2021). Since EM should not suffer from the same technical limitations, it seems likely that these states are indeed short-lived.

Generally, completely flat clathrin coats are rare in our data. Instead, we observed that clathrin coats quickly acquire curvature already early in endocytosis. This agrees with the early observation made using EM (Heuser, 1980) that even flat structures already had some small degree of curvature. Interestingly, an enrichment of pentagons was observed towards the edges of flat lattices, leading to the hypothesis that even without additional formation of pentagons these flat lattices have the built-in capacity to become curved, and are not very stable. This notion is in agreement with recent demonstrations that flat clathrin coats store energy, which can drive coat bending without additional factors (Sochacki et al., 2021; Tagiltsev et al., 2021).

The importance of curvature generation for clathrin coat maturation is supported by previous studies that suggest failure to initiate significant curvature as a hallmark of abortive events (Loerke et al., 2009; Wang et al., 2020). Observed in live-cell studies, these dim and short-lived events fail to significantly increase in intensity and do not undergo scission (Loerke et al., 2009; Mettlen et al., 2009). As structures in our data set mostly contain at least shallow curvatures at minimally half the final coat area, we are likely not capturing these abortive events, and they have negligible impact on our analysis.

### The cooperative curvature model

Here, we visualized the shapes of clathrin coats with high 3D resolution. This was crucial to precisely measure the clathrin curvature throughout the endocytic timeline ranging from flat to highly curved structures. Especially shallow curvatures are hard to accurately assess with 2D imaging. Thus, our data enabled us to robustly test different growth models for clathrin during endocytosis.

The constant curvature model predicts a continuous increase in surface area, constant coat curvature, as well as a rim length that increases until a half spherical geometry is reached, and then decreases again. Our experimental data, most notably the increase in curvature and monotonical decrease in rim length, are incompatible with these predictions, thus ruling out the constant curvature model (Figure 3, green lines).

The constant area model, in a strict interpretation, on the other hand predicts the coat to assemble completely on a flat membrane, after which its area stays constant and only curvature increases to form a spherical vesicle. These predictions agree reasonably well with the data for curvature propagation during CME, but fail to explain the monotonic increase in coat surface over time. Thus, the continuous increase of surface area that we observed here rule out the constant area model as well (Figure 3, blue lines). Interestingly, earlier work compatible with the constant area model already suggested the presence of shallow curvatures in even the flattest clathrin coated structures (Avinoam et al., 2015; Bucher et al., 2018; Heuser, 1980)

We found that around half of the clathrin coat area has preassembled when plasma membrane invagination begins (Figure 2D, 4C), agreeing well with previous reports (Bucher et al., 2018; Scott et al., 2018), after which the coat keeps growing gradually. Based on this observation, we developed a mathematical model that for the first time considers that curvature is generated in a positive (non-linear) feedback loop. Our cooperative curvature model assumes that net area growth of the clathrin coat occurs via triskelia addition to the many available binding sites at the coat edge. We proposed that this process depends on the number of free binding sites, which scale with the rim length, and can be described by a single kinetic constant *k*_on_. Triskelia addition within the coat is still likely to occur at a lower rate, as clathrin lattices can be unsaturated and have defects (Frey et al., 2020; Sochacki et al., 2021) and triskelia can exchange (Avinoam et al., 2015), which is energetically favorable for curvature changes within these lattices (Frey and Schwarz, 2020; Sochacki et al., 2021; Tagiltsev et al., 2021). While growth at the rim accounts for increase in surface area, curvature is most likely generated in the bulk of the coat. Here we assumed a non-linear relation between the rate of curvature increase and curvature, which reflects cooperativity in the lattice, e.g. due to rearrangements of neighboring triskelia or larger regions thereof. This assumption of cooperativity is supported by recent experiments, which suggest clathrin to exhibit curvature sensing properties, preferentially assembling at pre-bent membranes (Sochacki et al., 2021; Zeno et al., 2021). The model described above shows an excellent fit between theory and experiment, and further predicts a square root dependence of theta over time. This describes curvature generation during clathrin coat maturation as a non-linear mechanism driven by a positive feedback of multiple triskelia, slowing down once the coat approaches a preferred degree of curvature.

Even though our model agrees very well with the data, it still exhibits certain limitations worth mentioning:

Firstly, it only considers the coat itself and ignores the plethora of other proteins within the endocytic machinery. These include numerous BAR domain proteins that are dynamically recruited and disassembled during endocytosis and thus are bound to influence membrane curvature at endocytic sites, as well as a variety of clathrin adaptor proteins, whose presence or absence could explain the cell-type specific differences in average vesicle sizes that we observed (Supplementary Figure 5). Taken together, these factors could explain the imperfection of our model in the very beginning, and the final part of the timeline (Figure 4A), where vesicle scission is driven by the fast-acting mechanoenzyme dynamin. Additionally, we only modeled clathrin recruitment and ignore clathrin disassembly, which could be mediated by adaptor unbinding (Taylor et al., 2011) and uncoating factors including auxilin (He et al., 2020) that are recruited before the end of vesicle scission. We also assumed that the clathrin coat has constant properties, most notably that the intrinsic coat-driven curvature generation towards its preferred curvature occurs unidirectionally and remains the same throughout endocytosis (equation 3). It is however likely that the properties of the clathrin coat change during endocytosis, e.g. by coat stiffening or increasingly tight interactions between triskelia (Frey and Schwarz, 2020; Frey et al., 2020; Sochacki et al., 2021).

Secondly, we reconstructed an average trajectory of clathrin coat remodeling, generated from many individual static snapshots, thereby averaging conceivable different pathways of CME. However, different pathways with substantially changed relationships between the parameters like curvature and theta would be visible in the corresponding plots as separate point clouds. This is not the case, rather we observed a continuous correlation between curvature and theta following a single trajectory, indicating that CME follows a single, stereotypic pathway. We did identify a small, disconnected population of sites in our data set that most likely originate from a distinct cellular mechanism (Supplementary Figure 3 and 4). While this indicates that our approach should capture potentially different pathways of clathrin coated vesicle formation, we cannot exclude that minor mechanistic variations are included into the average trajectory.

Clathrin recruitment has been quantified extensively using live-cell microscopy, resolving the sequential recruitment and dynamic characteristics of many important endocytic components (Aguet et al., 2013; Cocucci et al., 2012; Doyon et al., 2011; He et al., 2017; Jin et al., 2021; Loerke et al., 2009; Mettlen et al., 2009; Saffarian et al., 2009; Schöneberg et al., 2018; Taylor et al., 2011). We wondered if it is possible to correlate our pseudotime reconstruction with previously reported real-time dynamics of clathrin, which shows an initial fluorescence intensity increase, a plateau, and finally a sharp intensity decrease after vesicle scission. While we observed a correlation between the number of clathrin localizations and surface area (Supplementary Figure 7), we note that indirect immunolabeling is not always quantitative, which complicates a direct comparison of previously reported live-cell fluorescence intensity over time and our pseudotime trajectory data. Nevertheless, we speculate that the initial fast intensity increase in live-cell studies likely corresponds to the initial growth of the flat clathrin coat that escapes our detection due to the fast dynamics and small size. The slower fluorescence increase and subsequent plateauing then coincides with curvature generation and final addition of triskelia at the coat rim as resolved in detail in our pseudotime data. To better understand the correlation between changes in the nanoscale architecture of clathrin coats and dynamic consequences, we would ultimately require a method that combines high structural with temporal resolution. In a recent publication, this was attempted using a live-cell 2D superresolution microscopy approach (Willy et al., 2021). This study reported an increasing projected area of clathrin over time, suggested that curvature is present in the earliest detectable clathrin structures, and concluded that CME follows the constant curvature model. Although we also find that completely flat structures are rare and curvature is initiated before the final surface is acquired, our data is entirely incompatible with the CCM. This is especially true in the first half of endocytosis with shallow but progressively increasing curvatures (Figure 3), which is challenging to measure using 2D imaging. This shows that it remains highly desirable to ultimately image the 3D nanoscale architecture of the clathrin coat in living cells in real time.

In summary, we characterized the dynamics and geometries of the clathrin coat in endocytosis by combining 3D superresolution microscopy, quantitative analysis, and mathematical modeling. We found that clathrin bending and assembly occur simultaneously along a precisely defined trajectory. We anticipate that this work will be foundational to further study the structural mechanism of endocytosis, both under physiological conditions, and in disease, where changes in CME likely occur very frequently and have just recently been shown to have profound functional consequences (Moulay et al., 2020; Xiao Guan-Yu et al., 2018).

## Acknowledgements

We thank the entire Ries and Kaksonen labs for fruitful discussions and support. This work was supported by the European Research Council (ERC CoG-724489 to J.R.), the National Institutes of Health Common Fund 4D Nucleome Program (Grant U01 to J.R.), the Human Frontier Science Program (RGY0065/2017 to J.R.), the EMBL Interdisciplinary Postdoc Programme (EIPOD) under Marie Curie Actions COFUND (Grant 229597 to O.A.), the European Molecular Biology Laboratory (M.M., A.T., Y.-L. W. and J.R.) and the Swiss National Science Foundation (grant 310030B_182825 and NCCR Chemical Biology to MK). O.A is an incumbent of the Miriam Berman Presidential Development Chair.

## Author contributions

M.M., O.A., M.K. and J.R. conceived the study, M.M., A.T., J.M. and O.A. performed experiments, M.M., A.T., Y.-L.W., J.M., O.A. and J.R. analyzed superresolution data; F.F. and U.S. developed the cooperative curvature model; J.R. supervised the study. M.M., A.T., F.F., J.R. wrote the manuscript with input from all authors.

## Material and methods

### Cell culture

SK-MEL-2 cells (gift from David Drubin, UC Berkeley, described in (Doyon et al., 2011)) were cultured adherently as described previously (Li et al., 2018) in DMEM F12 (Dulbecco’s modified Eagle’s medium with Nutrient Mixture F-12) with GlutaMAX, phenol red (Thermo Fisher; 10565018), 10% (v/v) FBS, ZellShield (Biochrom AG, Berlin, Germany), at 37 °C, under an atmosphere with 5% CO_2_ and 100% humidity.

U2OS cells (Cell Line Services; #300174) were cultured adherently as described previously (Thevathasan et al., 2019) in high-glucose DMEM supplemented with 10% FBS, 2 mM l-glutamine, non-essential amino acids, ZellShield at 37 °C under an atmosphere with 5% CO_2_ and 100% humidity.

3T3 mouse fibroblasts (gift from Alba Diz-Muñoz, EMBL Heidelberg) were cultured adherently in DMEM (4.5 g/L D-Glucose) supplemented with 1× MEM NEAA (catalog no. 11140-035, Gibco), 1× GlutaMAX (catalog no. 35050-038, Gibco) and 10% (v/v) fetal bovine serum (catalog no. 10270-106, Gibco) at 37°C under an atmosphere with 5% CO_2_, 100% humidity.

### Sample preparation for superresolution imaging of clathrin-coated pits

Cells were fixed as described previously (Li et al., 2018) using 3% (w/v) formaldehyde 10 mM MES, pH 6.1, 150 mM NaCl, 5 mM EGTA, 5 mM d-glucose, 5 mM MgCl_2_ for 20 min. Fixation was quenched in 0.1% (w/v) NaBH4 for 7 min. The sample was washed three times with PBS and permeabilized for 15 min with 0.01% (w/v) digitonin (Sigma-Aldrich, St. Louis, MO, USA) in PBS. The sample was then washed twice with PBS and blocked for 1h with 2% (w/v) BSA/PBS, washed with PBS, and stained for 3-12 h with anti-clathrin light chain (sc-28276; Santa Cruz Biotechnology, Dallas, TX, USA) and anti-clathrin heavy chain rabbit polyclonal antibodies (ab21679; Abcam, Cambridge, UK) in 1% (w/v) BSA/PBS. After three washes with PBS the sample was incubated for 3-4h with a secondary donkey anti-rabbit antibody (711-005-152; Jackson ImmunoResearch, West Grove, PA, USA) that was conjugated to Alexa Fluor 647-NHS at an average degree of labeling of 1.5. The sample was then washed three times and mounted for imaging in blinking buffer (50 mM Tris/HCl pH 8, 10 mM NaCl, 10% (w/v) D-glucose, 500 μg ml-^1^ glucose oxidase, 40 μg ml-^1^ glucose catalase, 35 mM MEA in H2O).

For the analysis of the disconnected population of sites, 3T3 cells were transfected with a plasmid encoding the sigma 2 subunit, fused to GFP (a kind gift from Steeve Boulant, University of Florida), to obtain cells transiently expressing AP2-GFP. The transfection was performed using a Lipofectamine™ 2000 reagent (Life Technologies) according to the manufacturer’s recommendations: 1 μg DNA was mixed with 50 μL OptiMEM I (ThermoFisher). The same was done for 3 μL Lipofectamin in 50 μL OptiMEM I. Both solutions were incubated for 5 min and then mixed together and incubated for additional 10 min, at room temperature. The media of previously seeded cells was exchanged to pre-warmed OptiMEM I, to which the DNA-Lipofectamin solution (100 μL) was dropwise added. After approximately 24 hours of incubation (at 5% CO_2_, 37 °C), the medium was exchanged with fresh growth medium. After additional incubation for approximately 16 hours, cells were fixed according to the same protocol described above.

### Superresolution microscopy

SMLM images were acquired at room temperature (24 °C) on a customized microscope (Mund et al., 2018) with a 160x NA 1.43 oil-immersion objective (Leica, Wetzlar, Germany). Illumination was done using a a LightHub laser combiner (Omicron-Laserage Laserprodukte, Dudenhofen, Germany) with Luxx 405, 488, and 638 Cobolt 561 lasers, which were triggered using a Mojo FPGA (Embedded Micro, Denver, CO, USA) for microsecond pulsing control of lasers. The lasers were guided through a speckle reducer (LSR-3005-17S-VIS; Optotune, Dietikon, Switzerland) and coupled into a multimode fiber (M105L02S-A; Thorlabs, Newton, NJ, USA). The output of the fiber is magnified and imaged into the sample. Fiber generated fluorescence was removed using a laser clean-up filter (390/482/563/640 HC Quad; AHF, Tübingen, Germany). The focus was stabilized using a closed-loop system, based on reflecting the signal of a near-infrared laser by the coverslip and detecting the resulting signal on a quadrant photodiode, resulting in focus stabilities of ±10 nm over several hours. Fluorescence emission was filtered using a 676/37 or a 700/100 bandpass filter (AHF) and recorded by an EMCCD camera (Evolve512D; Photometrics, Tucson, AZ, USA). Typically, 100,000-300,000 frames were acquired using 15-ms or 30 ms exposure times and laser power densities of ~15 kW/cm^2^. 405-nm laser intensity was adjusted automatically by changing pulse duration in order to keep the number of localizations per frame constant during the acquisition.

For the analysis of the disconnected population of sites, one diffraction limited image was additionally acquired before SMLM imaging. For this, a 488 laser at 1.4 kW/cm^2^ was used to take a single frame at 30 ms exposure time. Emission was filtered using a 525/50 bandpass filter (AHF).

### Data analysis

All data analysis was conducted in SMAP ((Ries, 2020) based on MATLAB and available as open source under github.com/jries/SMAP).

#### Superresolution image reconstruction

For fitting the localizations, peaks were detected as maxima in raw camera images after applying a difference-of-Gaussian filter. At those positions, cropped images of 13×13 pixels were fitted with an experimentally derived PSF model (free fitting parameters: *x*, *y*, *z*, photons per localization, background per pixel), using an MLE (Maximum likelihood estimation) fitter (Li et al., 2018). The *x*, *y* and *z* positions were corrected for residual drift by a custom algorithm based on redundant cross-correlation. In short, the data was distributed into 10 time bins of equal length. For each bin a superresolution image was reconstructed. We then calculated the image cross-correlations among all superresolution images and extracted the relative displacements in x and y were from the position of the maximum in the cross-correlation images. We then calculated the drift trajectory that best describes the relative displacements. In a second step, the z-drift was measured in an analogous way, using intensity profiles in z instead of images. Localizations persistent over consecutive frames (detected within 35 nm from one another and with a maximum gap of one dark frame) were merged into one localization by calculating the weighted average of *x*, *y* and *z* positions and the sums of photons per localization and background. Localizations were filtered by the localization precision in x,y (0-20 nm) and z (0-30 nm) to exclude dim localizations. The modal value for the localization precision *σ* was 3.9 nm in x/y and 12.5 nm in z, leading to a resolution estimate (calculated using the Full Width Half Maximum (FWHM) using FWHM =2√ (2/π2)σ) of 9.2 nm in x/y and 29.4 nm in z (typical values based on representative image). Superresolution images were constructed with every localization rendered as a two-dimensional spherical Gaussian with a width of 3 nm. The red hot color map used represents the density of localizations, and is scaled in a way that 0.03% of the pixel values are saturated.

#### Quantitative geometric analysis of clathrin-coated structures

Clathrin-coated structures were segmented semi-automatically. First, we manually defined a region of interest excluding the edges of the cell. Then, the image was blurred using a Gaussian filter with a sigma of 100 nm, and peaks were detected using a manually set threshold. This typically yielded several hundreds of sites in a region of 30×30 μm. These candidate sites were curated manually, and only single, well-isolated clathrin-coated structures were retained in the dataset.

Next, these structures were analyzed using LocMoFit, an MLE-based model fitting framework that we developed recently (Wu et al., 2021). LocMoFit directly fits localization coordinates with the probability density function (PDF) of a parametrized geometric model. In this study, we modeled clathrin-coated structures with a hollow spherical cap parameterized by the surface area *A* and the closing angle *θ*. *θ* is defined as the angle between the two vectors that point to the pole and to the rim, respectively, from the center of the sphere. The position of the model is defined as the center of mass of the cap. In practice, we discretized the cap by placing spiral points (Saff and Kuijlaars, 1997), or the spherical Fibonacci lattice, on the surface of the cap to approximate an even distribution. In LocMoFit, these points were treated as discrete fluorophore coordinates when constructing the PDF of the model. During fitting, additional parameters including the center position and the orientation of the model were determined with respect to the fluorophore coordinates, and an extra uncertainty and the background weight were applied to the PDF. After fitting, the sphere radius is derived as 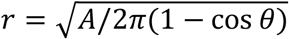, projected area as *A_p_* = *π* sin^2^ *θ* and edge length as *ε* = 2*π*sin *θ*. For some flat sites where the fit produced slightly negative curvature values, curvature *H* and *θ* were set to 0 nm^-1^ and 0° respectively in order to approximate them as completely flat.

After model fitting, a second curation step was performed. With this we ensured that only well fitted sites are included in the final data set. Sites were excluded if they were upside down (clearly originating from an upper membrane), double-sites with two sites clearly connected to each other or an adjacent flat structure, large plaques or structures non-distinguishable from an antibody cluster.

#### Pseudo-temporal reconstruction of clathrin remodeling during endocytosis

To sort endocytic sites in pseudotime, they were sorted by the closing angle *θ*, assigning each site a rank index. Flat sites with a manually assigned H = 0 nm^-1^ and *θ* = 0 were all assigned an index of 0. As pseudo-temporal sorting assumes that all sites are part of the same endocytic trajectory, endocytic sites with curvatures above a cell line specific threshold (*H* > 0.016 nm^-1^ for SK-MEL-2; *H* > 0.013 nm^-1^ for U2OS; *H* > 0.014 nm^-1^ for 3T3) that form a visibly disconnected point cloud (Supplementary Figures 3 and 4) were excluded for this analysis (Supplementary Figure 5). To compute pseudo-temporal averages, the sites were spatially and rotationally aligned, and rescaled by the average radius of all sites within the respective bin. Because the geometric model is rotationally symmetric, we then performed rotational averaging by generating 72 duplicates of each structure, rotating them by 5 degrees with respect to each other, and averaging them. The supplementary movie was computed using a sliding window approach, where each frame shows the average of 30 sites, and the frame-to-frame increment is 20 sites. The median pseudotime of those 30 sites is indicated.

#### Further data analysis

All data that resulted from the quantitative geometric description of clathrin-coated structures were further analyzed in R (R Core Team, 2020). Fitting of the growth models (Tables 1-3) was performed on a filtered data set, excluding disconnected sites above a cell line specific threshold, and sites of negative curvature. During fitting, for sites with *θ* = 0° we set *θ* = 0.0001° to avoid division by 0.

For depicting the growth models in Figure 3 and 4, as well as Supplementary Figure 5, parameters resulting from the *H*(*θ*) fit were used and mapped to the *A*, Ɛ and *A_p_* data.

#### Simulations

Simulations were performed using the simulation engine for SMLM data implemented in SMAP and LocMoFit as described in (Thevathasan et al., 2019). The realistic simulations were based on a two-state (bright and dark) fluorophore model plus bleaching (Sage et al., 2019) and parameters (number of photons, background photons, and fluorophore on-time *t*_1_) extracted from our experiment. (1) First, we defined an equally distributed closing angle *θ* from 0 to 180° and calculated the surface area *A*(*θ*) = 2*π*(1 – cos*θ*)/*H*(*θ*)^2^ (see Supplementary Note for details). Here *H*(*θ*) is defined as in Equation 3, with fitting parameters determined in SK-MEL-2 (see Table 1). (2) With the defined model parameters, we generated protein positions for each simulated structure by taking randomly drawn *N* α *A* samples from the PDF of the hollow spherical cap with no uncertainty. (3) With a probability *p*_label_ = 0.6, a fluorescent label was created at a protein position. (4) Linkage displacements in x, y and z were added to a label and were determined as normally distributed random variables with a variance corresponding to the linkage error of 5 nm. The fluorophore is assumed to be freely rotating three-dimensionally between different blinks. (5) Each fluorophore appeared at a random time and lived for a time *t_l_*, determined as a random variable from an exponential distribution with a mean of 1.6 frames. (6) A label had a probability *p*_react_ = .5 to be reactivated and then appeared at a random later time point, otherwise it was bleached. (7) When it was on, a fluorophore had a constant brightness. Thus, the brightness in each frame was proportional to the fraction of the time the fluorophore was on in each frame. (8) The emitted photons in each frame were determined as a random Poisson variable with a mean value corresponding to the average brightness in the frame. (9) For each frame, we calculated the CRLB (Cramér-Rao lower bound) in x, y and z from the number of photons (with a mean of 11,000) and the background photons (130 per pixel) based on the theoretical Gaussian PSF (Mortensen et al., 2010) or a 3D cspline PSF model derived from beads calibrations (Li et al., 2018). This error was added to the true x, y and z positions of the fluorophores as normally distributed random values with a variance corresponding to the respective calculated CRLB.

## Supplementary Information

**Supplementary Figure 1.**
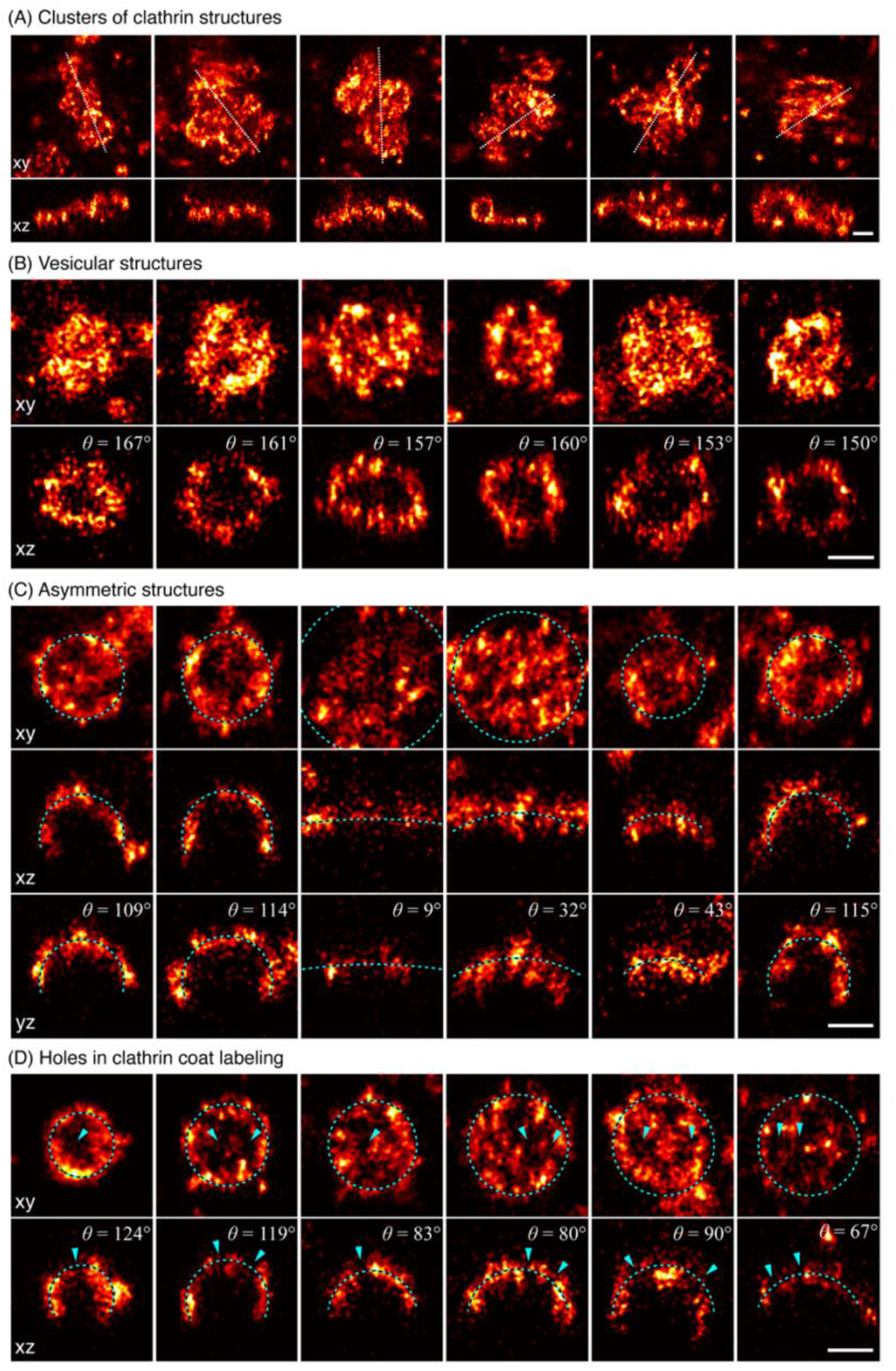
Examples of diverse clathrin coat structures. (A) Large clusters of clathrin molecules excluded from further analysis. Shown in top view (xy) with dotted line indicating a 50 nm-thick z-slice (xz) shown below. (B) Vesicular structures are sometimes fitted with a lower θ as expected. (C) While a spherical model describes structure of most endocytic clathrin coats faithfully, there are few cases, as exemplified here, where the elliptical and irregular shape of an assembling coat is difficult to approximate with a simple geometric model. Two orthogonal 50 nm-thick z-slices are shown here in xz and yz, and the respective spherical model fit is plotted as a dotted line. (D) Non-continuous labeling of clathrin manifests itself as holes in the coat, indicated with a blue arrow. All scale bars are 100 nm.

**Supplementary Figure 2.**
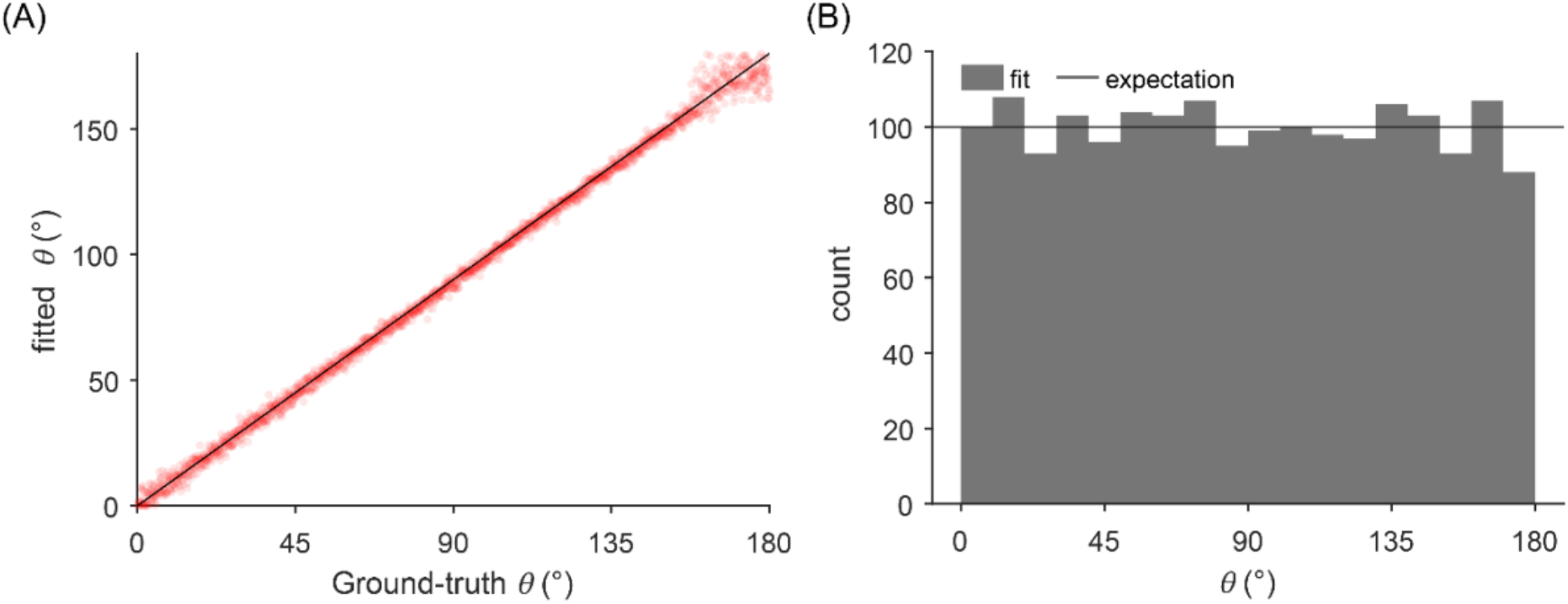
Estimation error of the closing angle θ. (A) The comparison of fitted θ of simulated clathrin coat structures to corresponding ground-truth θ shows no systematic bias and a narrow spread of the error across the ground-truth although the spread increases when θ < 20° or θ > 160°. We reasoned that in the earlier range, where the structures are flat, the slightly increased error corresponds to the insensitivity of θ to flat structures. In the later range, where the vesicles are almost closed, the error was caused by undisguisable tiny holes corresponding to the real vesicle openings and unlabeled clathrin in the coat. The fitted structures were simulated to have similar quality as the experimental data and to distribute evenly across θ. (B) The distribution of θ shows no significant error compared to the expectation corresponding to the evenly distributed θ, except for the small potential underestimation of entirely closed coats.

**Supplementary Figure 3.**
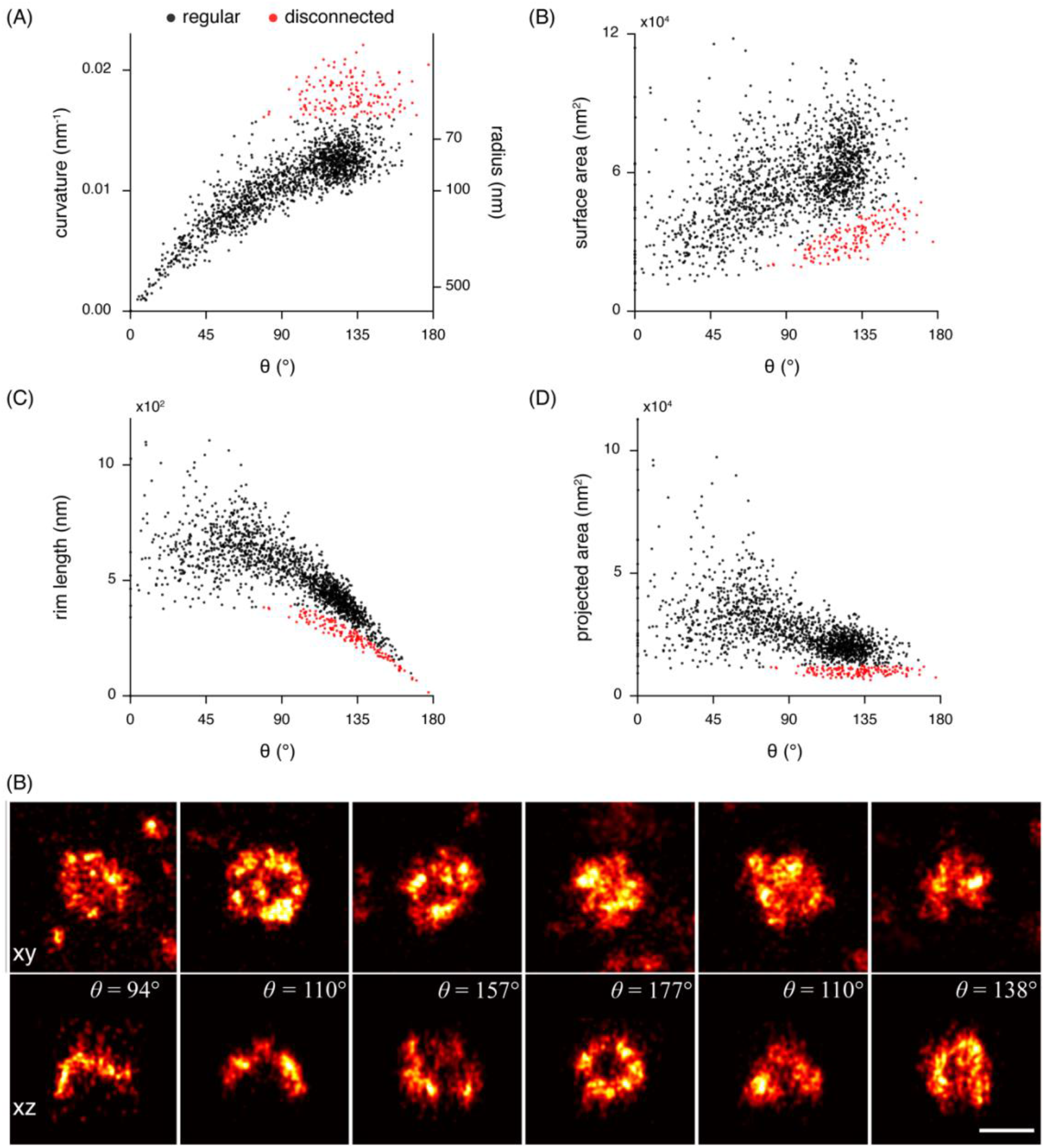
Non-endocytic clathrin structures. Results of LocMoFit analysis of clathrin structures in SK-MEL-2 cells (N = 13 cells, n = 1798 sites). (A) A strong correlation between curvature and θ can be observed for most structures (n = 1645 sites, black). A disconnected point cloud (n = 153 sites, 8.5%, red) indicates the presence of endocytosis-unrelated clathrin structures. The same distinct population of data points can be observed for the (B) surface area, (C) rim length, and (D) projected area. (B) Example structures from the disconnected population of sites in top view (xy) and 50 nm-thick z-slices (xz) and their respective θ values. Scale bar is 100 nm.

**Supplementary Figure 4.**
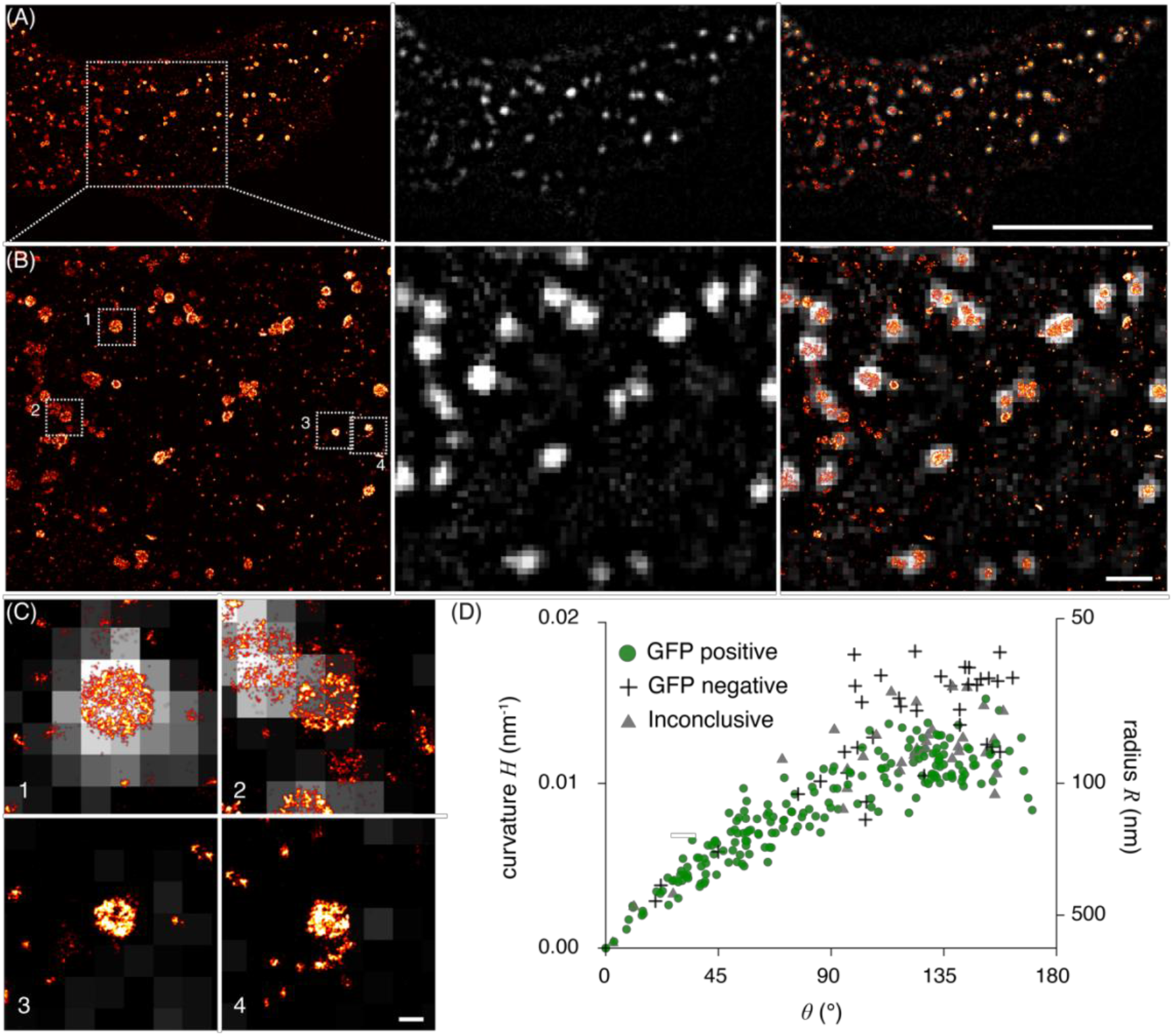
Analysis of clathrin coats not following general trajectory of curvature generation. (A) 3T3 cell transiently overexpressing AP2-GFP. (left) Single-molecule localization microscopy image of immunolabeled clathrin. (middle) Diffraction limited image of the AP2-GFP signal. (right) Overlay of the two targets (scale bar: 10 μm). (B) Enlarged image of the section indicated in (A) (scale bar: 1 μm). (C) Example sites indicated in (B) (scale bar: 100 nm). (1) Example for a structure annotated as “GFP positive”. (2) Example for an “inconclusive” GFP signal. (3 and 4) Example of “GFP negative” structures”. (D) Analysis results when estimating theta and curvature from clathrin structures, and annotating them depending on their AP2 signal. (N = 3 cells and n = 277 sites). No AP2-GFP positive structures are found in the disconnected population of sites, suggesting that they are most likely not generated via clathrin mediated endocytosis.

**Supplementary Figure 5.**
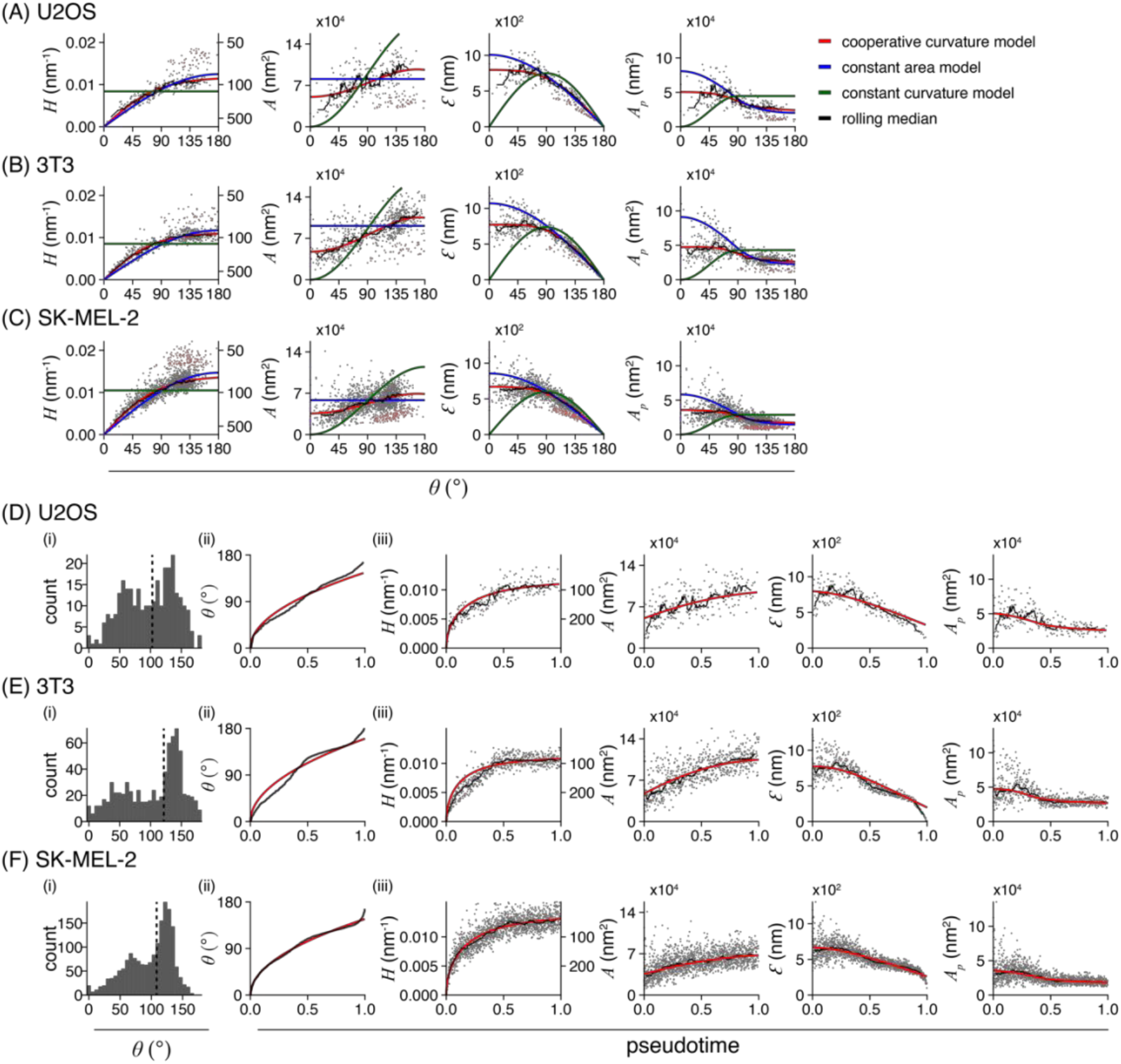
Clathrin coat remodeling in three different cell lines. (A-C) Results of LocMoFit analysis for clathrin structures. Different growth models are fitted to curvature H over θ. The resulting fitting parameters are then used to map the same models also over surface area, rim length, and projected area (left to right). Purple: completely flat sites with H = 0 nm^-1^ and θ = 0. Red: disconnected sites that were excluded from the fitting. Black line: rolling median (window width = 5% of total number of sites) (A) U2OS (N = 3 cell, n_grey_ = 241 sites, n_red_ = 53 disconnected sites, n_purple_ = 1 completely flat sites), (B) 3T3 mouse fibroblasts (N = 7 cells, n_grey_ = 688 sites, n_red_ = 51 disconnected sites, npurple = 8 completely flat sites), and (C) SK-MEL-2 cells (N = 13 cells, n_grey_ = 1631 sites, n_red_ = 153 disconnected sites, n_purple_ = 14 completely flat sites). (D-F) Temporal reconstruction of clathrin coat remodeling. (i) Distribution of θ slightly differs between cell lines, especially in the earlier states. Median θ shown as dotted lines correspond to 99.6° for U2OS; 121.4° for 3T3; and 108.5° for SK-MEL-2 cells. (ii) The cooperative curvature model (red line) highlights the square-root dependence between θ and pseudotime. (iii) The cooperative curvature model is used to describe the curvature H propagation over pseudotime. Resulting fitting parameters are then used to map the same model to surface area A, rim length Ɛ and projected area A_p_. A rolling median is plotted for in black (window width = 5% of total number of sites).

**Supplementary Figure 6.**
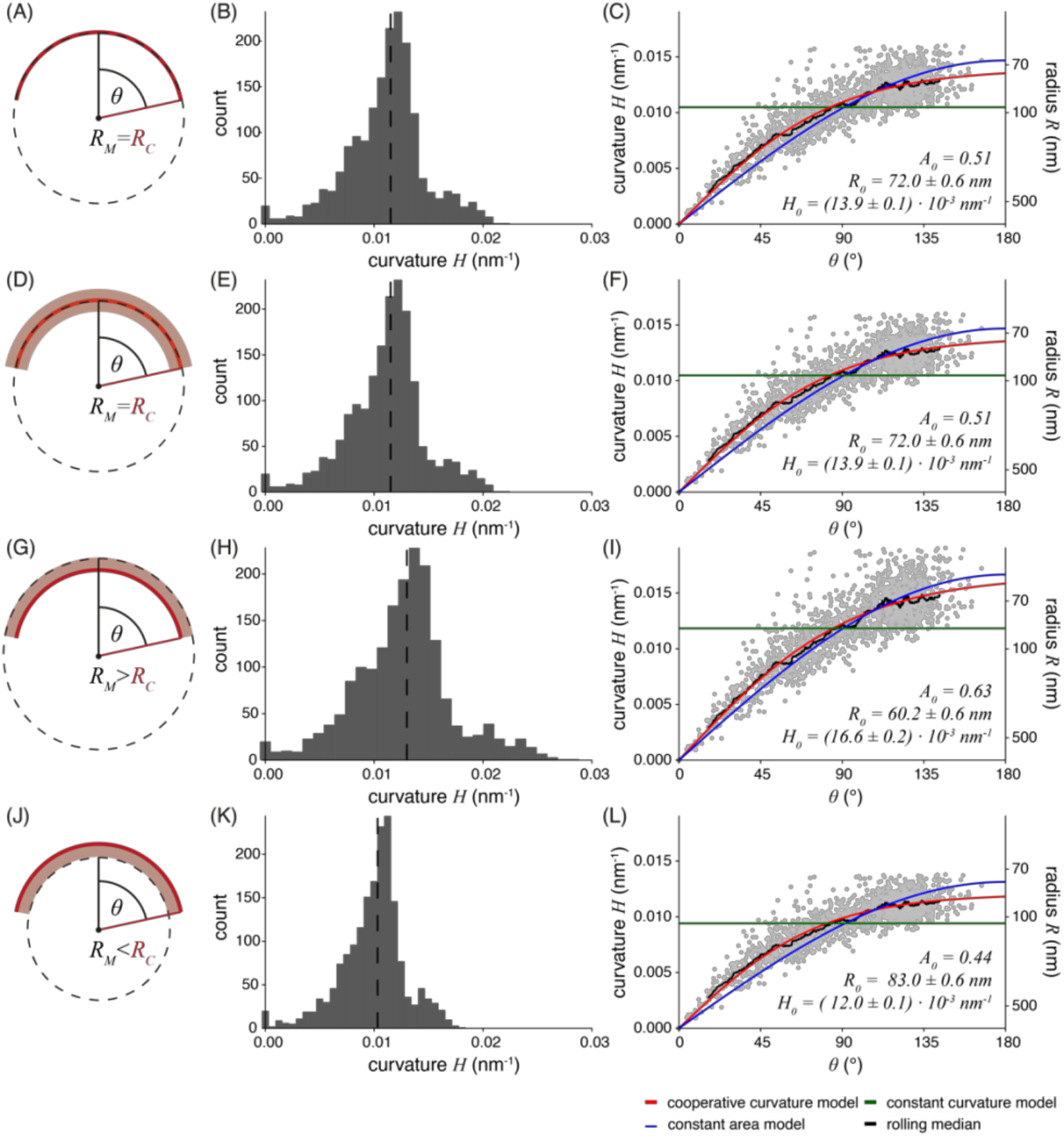
Impact of linkage error on geometric model fit. Indirect immunolabeling displaces the label from the target molecule by two antibodies. This generates a so-called linkage error of on average ±10 nm, and resulting localizations might not accurately represent the underlying structure of interest (Früh et al., 2021). (A) Ideal case of no linkage error, where the model (dotted black line) with radius R_M_ accurately represents the underlying clathrin coat (red) with a radius R_C_. (B) Histogram of quantified curvature with a median of 0.011 nm^-1^. (C) Models fitted to curvature propagation over theta θ. (D) Uniform displacement of localizations (light red, ± 10 nm) due to unbiased labelling by the antibodies. The radius R_M_ still accurately represents the true radius R_C_. (E) Histogram of the curvature, assuming no needed correction of R_M_. Median: 0.011 nm^-1^. (F) Models fitted to curvature propagation over theta θ remain the same as for (C). (G) Biased labeling of the antibodies (light red, + 10 nm) could result in an overestimation of R_M_ by 10 nm. (H) Histogram of the curvature corrected by subtracting 10 nm from quantified R_M_. Median: 0.013 nm^-1^. (I) Models fitted to corrected curvature propagation over theta θ. (J) Biased labeling could result in an underestimation of R_M_ by 10 nm. (H) Histogram of the curvature corrected for the overestimation in radius by adding 10 nm from quantified R_M_. Median: 0.010 nm^-1^. (I) Models fitted to corrected curvature propagation over theta. (C; F; I; L) Fitting parameters are A_0_: Fraction of surface area growing as a flat lattice before curvature initiation, defined as A_0_ = A(θ = 0.01)/A(θ = π); R_0_: preferred radius of the clathrin coat fitted with CoopCM; H_0_: Preferred curvature of the clathrin coat fitted with CoopCM. While the fitting parameters scale with the error in radius estimation, the relationships among the parameters and thus our mechanistic interpretation by the cooperative curvature model still holds true.

**Supplementary Figure 7.**
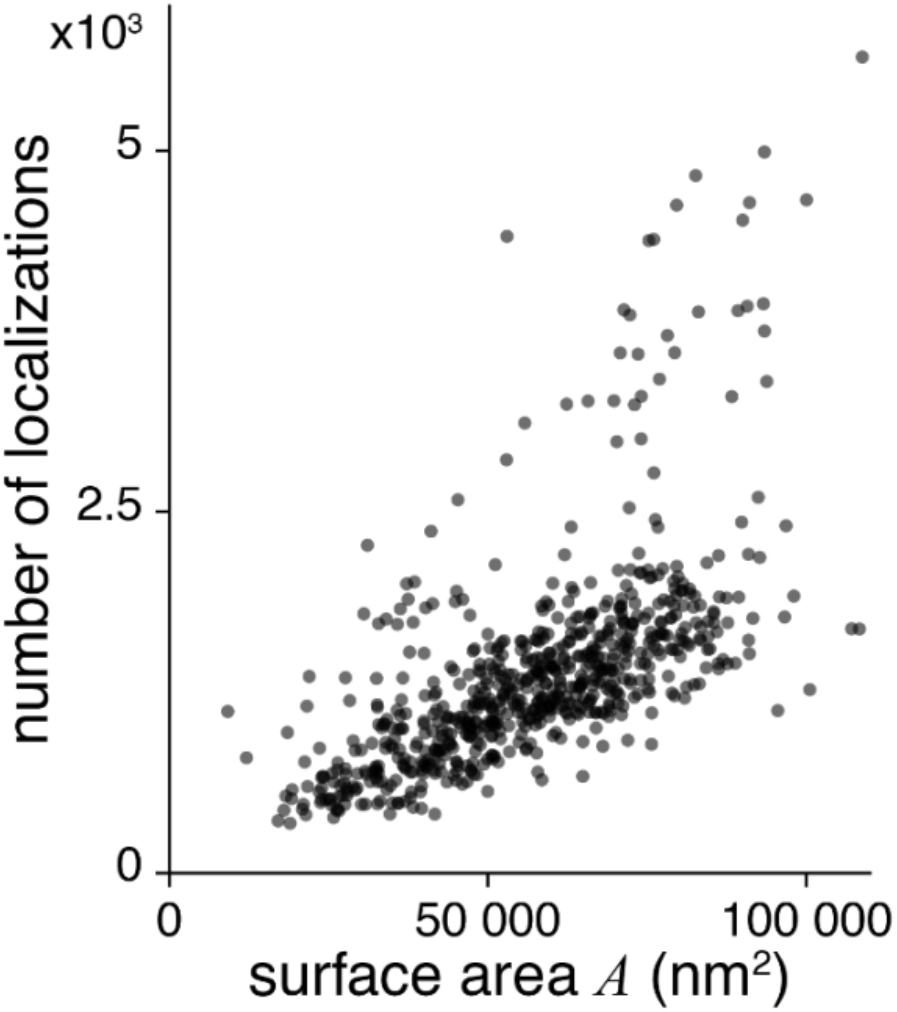
Number of localizations versus surface area. For N = 6 SK-MEL-2 cells, n = 700 sites the number of localizations found in one clathrin coated structure was extracted. This is plotted against the quantified surface area determined for each coat.

**Supplementary Figure 8.**
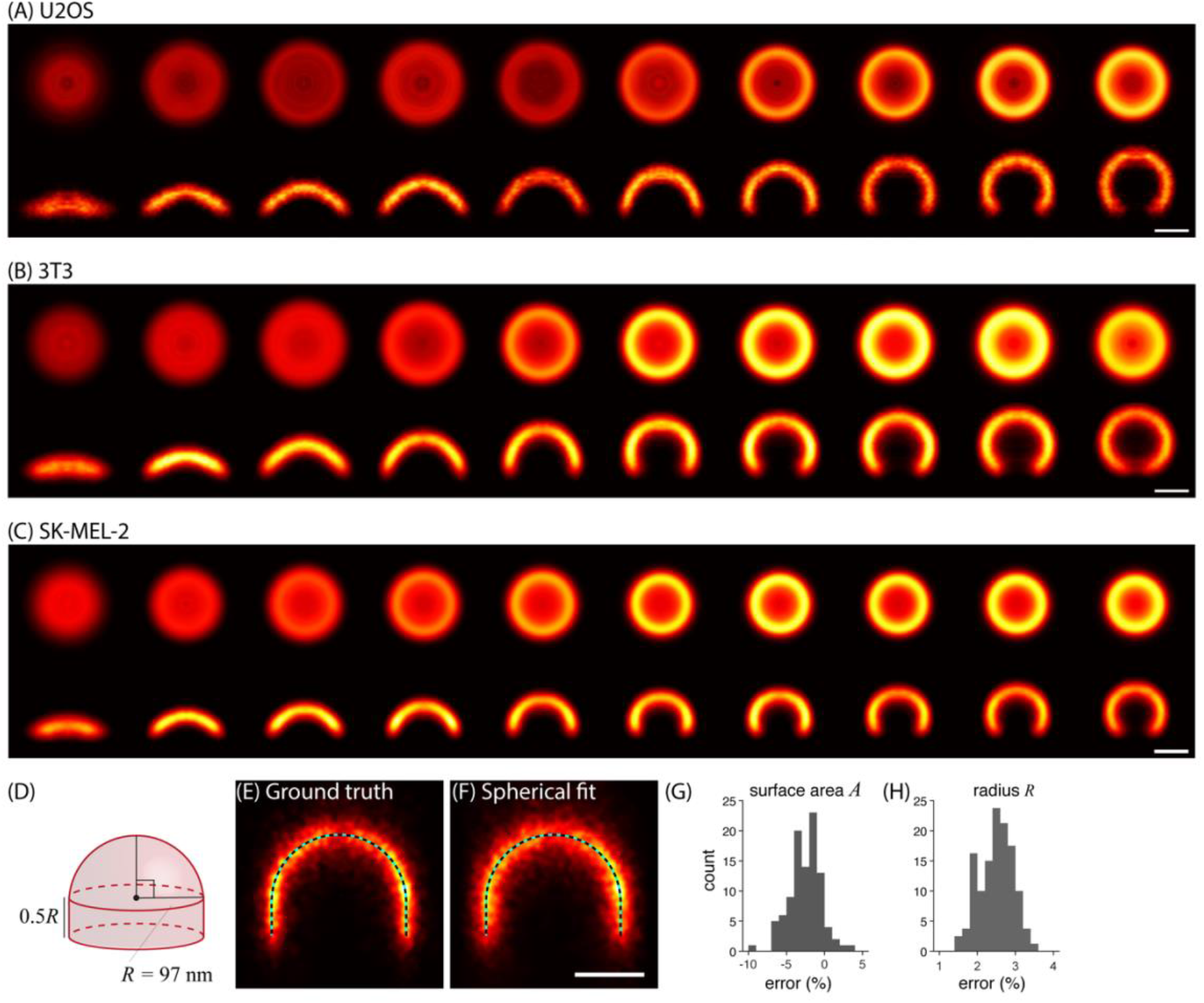
Averaging of clathrin coated structures. (A-C) Averages for distinct endocytic stages, resulting from all collected snapshots for (A) U2OS (n = 241 sites, N = 3 cells), (B) 3T3 (n = 688 sites, N = 7 cells), and (C) SK-MEL-2 cells (n = 1645 sites, N = 13 cells). Each bin contains the same number of snapshots of clathrin-coated structures sorted along their pseudotime (n_U2OS_ = 24 per bin, n_3T3_ = 68, n_SK-MEL-2_ = 164), so that all bins represent equally long pseudotime intervals. (D-H) Averaging preserves U-shapes in simulated endocytic sites. (D) A ground-truth structure was used for simulating U-shaped clathrin coats. The model for simulation was built by combining a hemisphere with a radius of R = 97 nm and a cylinder with the height D = 0.5 R. We choose the radius value according to the median radius of bin 6 in Figure 4G. This bin has a closing angle slightly larger than 90°. 20-nm thick cross-sections of the averages (N = 100 sites) registered based on (E) the ground truth and (F) the spherical fit are shown with the ground-truth U-shaped model (dotted line). Histograms of normalized estimation errors are shown for the parameters (G) surface area and (H) radius, with mean values of −2.6% and 2.5% respectively. The scale bar is 100 nm.

## Supplementary data

Supplementary data can be downloaded from https://oc.embl.de/index.php/s/sszVT1ZAUB2LOFW and will be uploaded to the BioStudies image data repository (https://www.ebi.ac.uk/biostudies/) upon publication.

### Supplementary movie (mp4 format, 1 file)

Pseudo-temporal movie of clathrin coat remodeling during endocytosis in SK-MEL-2 cells. Each frame corresponds to a sliding window average of 30 sites, with a frame-to-frame increment of 20 sites. The median pseudotime of those 30 sites is indicated.

### Localization data (csv format, 23 files)

Localization coordinates for all superresolution images, grouped by cell line (SK-MEL-2, 13 files; U2OS, 3 files; 3T3, 7 files).

The localizations were grouped (localizations that persisted over several adjacent frames and thus resulted from the same single fluorophore were combined into one localization) and filtered with a maximum localization precision of 20 nm in xy, and 30 nm in z. The first 2000 frames of each acquisition are excluded, since the blinking density is too high for reliable localization. Each file contains the columns *xnm* (x coordinate in nm); *ynm* (y coordinate in nm); *znm* (z coordinate in nm); *locprecnm* (localization precision in xy in nm); *locprecznm* (localization precision in z in nm); *frame; site_ID* (marks all localizations belonging to the respective segmented clathrin structure; localizations outside of analyzed sites are put to 0); *filenumber* (identifier for acquisition file, unique for each cell line).

Identifiers for acquisition files and clathrin-coated structures are kept consistent between all supplementary data.

### Model fit results (csv format, 23 files)

Results from LocMoFit analysis of individual clathrin coated structures, grouped by cell line.

Columns are *ID* (identifier for individual structures); *file_number* (identifier for each acquisition file); *cell_line*; *disconnected_sites* (true for those sites that are disconnected from the main data cloud, see Supplementary Figures 2, 3). Model fit parameters calculated for each structure using LocMoFit include theta θ; radius *R*; curvature *H*; surface_area *A*; rim_length *ε;* and projected_area *A_p_*. Further included in the table are the corrected values theta_corrected and curvature corrected, where each structure fitted with a negative radius was manually put to θ = 0.0001° and curvature = 0 nm^-1^.

### Site gallery (PDF format, 3 files)

Snapshots of all analyzed clathrin coated structure and their respective model fit results.

For each structure, shown are the top view (left), as well as 50 nm side-view cross-sections through the structure at angles 0°, 45°, 90° and 135° (left to right). Shown are site number, theta θ, surface area *A*, and radius *R*. Scale bar is 100 nm.

### Movie gallery (mp4 format, 49 files)

Rotating movies of the 3D SMLM data (left) and renderings of the corresponding models (right) of all clathrin-coated structures in SK-MEL-2 cells. Shown are site identifiers and model fit results (theta θ, surface area *A*, and radius *R*). Scale bar is 100 nm.

## Supplementary note

### I. COOPERATIVE CURVATURE MODEL

We derive a kinetic model for clathrin-mediated endocytosis that is based on minimal assumptions based on our experimental observations. We first assume that the area *A* of clathrin coats grows mainly by addition of new triskelia at the edges [1]. With a local growth rate *k*_on_ and an edge length *Ɛ*, we have the simple growth law

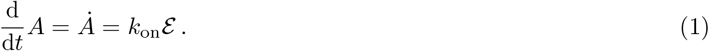

We also know from our experimental observations that all clathrin-coated pits have the geometry of a spherical cap with area *A*_cap_ = 2*πR*^2^(1 – *cos θ*), where *R* is the cap radius and *θ* the closing angle. The edge length then is *Ɛ*_cap_ = 2*πR* sin *θ*. Using these formulas on Eq. (1) we find

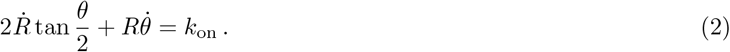

Because we parametrize the shape of the growing clathrin coat by the two dynamical variables *R*(*t*) and *θ*(*t*), but have only one growth equation up to now, we have to make additional assumptions to completely define our kinetic model.

As the main function of clathrin is to generate curvature, we now focus on coat curvature *H* = 1/*R* and assume that it increases with a basal rate *γ.* Experimentally we observe that clathrin patches first grow flat and then start to curve. Curvature generation in clathrin lattices is the combination of a preferred curvature of the single triskelion and cooperative effects in the lattice [2, 3]. Several mechanisms exist that might generate curvature during coat growth. First, clathrin triskelia are likely to be geometrically frustrated within the flat clathrin coat. To overcome this frustration, they would have to adjust their positions relative to their neighbors, such that their free energy is getting more favorable. Second, lattice vacancies could be filled up with new triskelia, as predicted theoretically [4]. Note that these vacancies would not necessarily relate to patch area, because the clathrin lattice consists of many overlapping arms. Filling of vacancies would however increase clathrin density and coat stiffness, and thus could also drive invagination of the coat [5]. We note that it was indeed confirmed experimentally that the clathrin density increases within the coat during invagination [2, 6]. Third, lattice pentagons, associated with lattice curvature, could form at the edge of the clathrin lattice and diffuse into the coat, similar to lattice defects on curved surfaces [7]. All three mechanisms could generate curvature with a certain rate which we assume to be constant during the initial stages of growth.

At late stages, increase of curvature has to stop and therefore we assume curvature saturation at a characteristic value *H*_0_, which is similar to, but different from the radius measured for clathrin cages. A simple estimate would be 40 nm for a typical radius of the membranes in the pits [8] plus 15 nm thickness of the clathrin coat [9]. These 55 nm would be larger than the 40 nm for clathrin cages, which are expected to be frustrated. If all triskelia in the lattice had achieved their optimal positions, a preferred or spontaneous curvature Ho should emerge. Thus a growth equation for *H* should have a stable fixed point generated by a higher order term. Since the corresponding mechanism is known to be of cooperative nature, larger groups of triskelia should be involved and the mechanism should start to dominate relatively late in the process, but then dominate quickly. Here we assume that the mechanism is proportional to *H*^2^ (rather than to *H*, with a linear law corresponding to a non-cooperative effect that starts to show early). Assuming that curvature generation is a geometrical effect, its time development should depend on the closing angle and we get

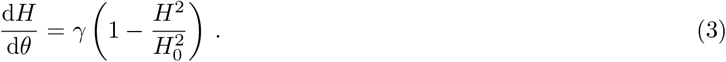

The stable fixed point at *H* = *H*_0_ can then be interpreted as corresponding to the preferred curvature of the mature coat.

Eq. (3) can be solved with the initial condition *H*(*θ* = 0) = 0 by

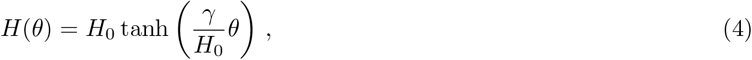

**TABLE I.** Coat curvature, coat area, and edge length summarized for the CCM, the CAM, and the CoopCM.

|  | CCM | CAM | CoopCM |
| --- | --- | --- | --- |
| $H$ | $1/R_{\text{CCM}}$ | $\sqrt{\frac{2\pi(1-\cos\theta)}{A_{\text{CAM}}}}$ | $1/R_{\text{coat}}$ |
| $A$ | $2\pi R_{\text{CCM}}^2(1-\cos\theta)$ | $A_{\text{CAM}}$ | $2\pi R_{\text{coat}}^2(1-\cos\theta)$ |
| $\mathcal{E}$ | $2\pi R_{\text{CCM}} \sin\theta$ | $\sqrt{\frac{2\pi A_{\text{CAM}}}{(1-\cos\theta)}} \sin\theta$ | $2\pi R_{\text{coat}} \sin\theta$ |

and defines the coat radius as a function of the closing angle

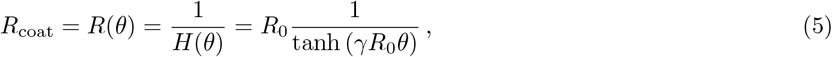

with *R*_0_ = 1/*H*_o_. We note that the expansion of Eq. (5) gives *R* ∝ 1/*θ* in leading order around the flat state.

We can combine Eq. (2) and Eq. (3) to find the full dynamics of coat invagination. Therefore, we compute the derivative of *R*_coat_

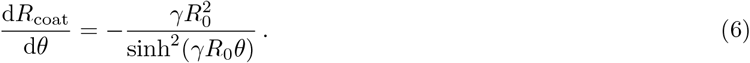

We use Eq. (5) and Eq. (6) and the chain rule *dR/dt* = *dR*/d*θ* d*θ*/d*t* to rewrite Eq. (2). Solving for 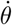 yields

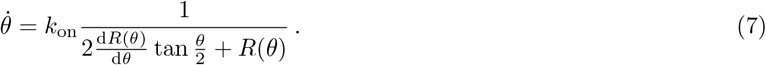

After expanding Eq. (7) up to leading order in *θ* we find

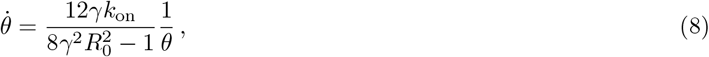

which is solved with the initial condition *θ*(*t* = 0) = 0 by

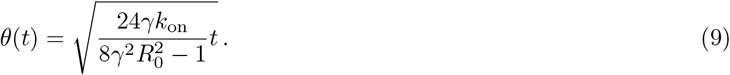

Thus the coat initially is flat (*θ* = 0) and then starts to generate curvature, increasing the closing angle with a square root dependence in time, which reflects the slowing down of curvature generation as preferred curvature is approached. We refer to this model as the cooperative curvature model (CoopCM) due to the assumption of a non-linear growth law.

### II. FIT RESULTS

To contrast the CoopCM with existing models, we fit it to experimental data for *H*(*θ*), *A*(*θ*) and *Ɛ*(*θ*). We compare the fitted curves to the constant curvature model (CCM), defined through the constant coat curvature 1/*R*_CCM_, and to the constant area model (CAM), defined through the constant coat area *A*_CAM_. The equations for the curvature *H*(*θ*) = 1/*R*(*θ*), the area *A*(*θ*), and the edge length *Ɛ*(*θ*) are summarized in Table I for all three models. In order to fit, we first filter the data according to curvature H, as described in the methods section of the main text. We then fit the different models to each of the data sets separately. For the CCM and CAM we get values for one parameter each, namely *R*_CCM_ and *A*_CAM_, respectively. For the CoopCM we obtain two parameter values *γ* and *H*_0_. For the CoopCM we also determine the area when invagination occurs *A*_0_ = *A*(*θ* = 0.01)/*A*(*θ* = *π*), which is the relative transition size where the flat-to-curved-transition occurs. By definition, invagination in the CCM starts simultaneously with coat assembly (*A*_0_ = 0), while invagination in the CAM starts at the maximum area (*A*_0_ = 1). Moreover, for the CoopCM we also compute *R*_0_. For all parameters we also determine the relative fit errors, given by *σ_x_/x*, where *σ_x_* is the standard deviation of the parameter *x* and *x* is the value of the parameter. This procedure allows us to compare the fit results to each other. The fitted parameter values for the different models and cell lines are summarized in the Supplementary Tables 1-3.

We then fit Eq. (9) to the data, describing *θ*(*t*). Using the values of *γ* and *H*_0_, we determine *k*_on_ from the fit. The resulting values, measured in units of the pseudo time 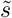, are summarized in the Supplementary Tables 1-3. We do not show the fit errors of *k*_on_, because they are small and result from the same fit of *θ*(*t*) and the errors of *γ* and *H*_o_, which does not allow us to discriminate between the different fits.

**TABLE II.** BIC for the curvature fits *H*(*θ*) of the CCM, CAM and CoopCM for the analyzed cell lines.

|  | CCM | CAM | CoopCM |
| --- | --- | --- | --- |
| SK-MEL-2 cells | -19079 | -21218 | -21670 |
| c3T3 cells | -8119 | -9179 | -9498 |
| U2OS cells | -2784 | -3097 | -3190 |

### III. BAYESIAN INFORMATION CRITERION

Apparently, the CoopCM agrees better with the data compared to the CCM and the CAM. However, while the CCM and the CAM both only include one parameter, the CoopCM has two parameters. In order to base model selection on a statistical criterion and to account for the additional parameter of the CoopCM, we compute the Bayesian information criterion (BIC). The BIC takes into account both how well a model fits the data but also how many free parameters it includes to avoid overfitting. The BIC can be expressed by [10]

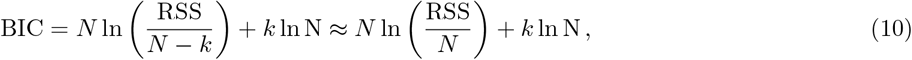

where we used that *N* ≫ *k* in the last step. Here RSS is the residual sum of squares 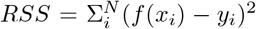 with *x*, and *y*, the measured data and *f* (*x_i_*) the fitted data, *N* is the number of data points and *k* is the number of free parameters. In general, when comparing different models, the model with the lowest BIC is preferred. In Table II the BIC is calculated from Eq. (10) for the curvature fits *H*(*θ*) for the different cell lines. For all cell lines, the BIC for the CoopCM is lowest. Therefore, we conclude that the additional parameter of the CoopCM is justified and that the CoopCM agrees best with the data.

### IV. LINEAR CURVATURE MODEL

As an alternative to the CoopCM model we now consider a linear curvature model. As in the CoopCM model, we assume that the coat curvature *H* = 1/*R* increases at a basal rate γ. In contrast to the CoopCM model however, we now assume that the increase in curvature at late stages is slowed down proportional to H and must stop at a value *H*_0_

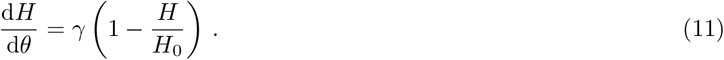

Eq. (11) can be solved by

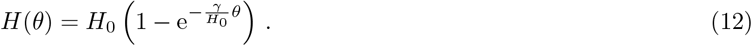

Eq. (12) also defines the coat radius *R* = 1/*H* as a function of the invagination angle *θ*

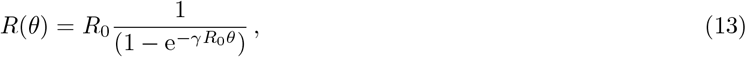

with *R*_0_ = 1/*H*_0_. The derivative of *R*(*θ*) reads

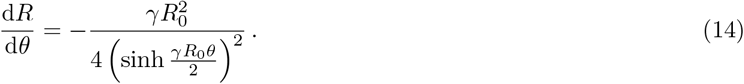

Now we use Eq. (13) and Eq. (14) on Eq. (7). After expanding up to second order in *θ* we find

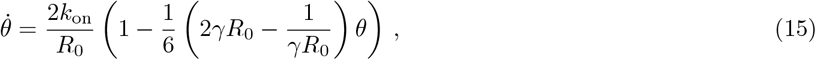

which is solved by

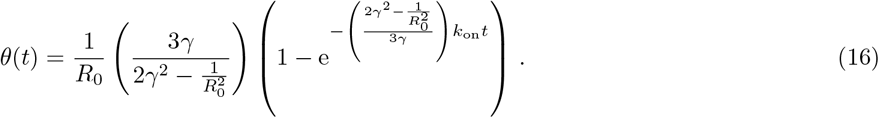

**Supplementary Figure 9.**
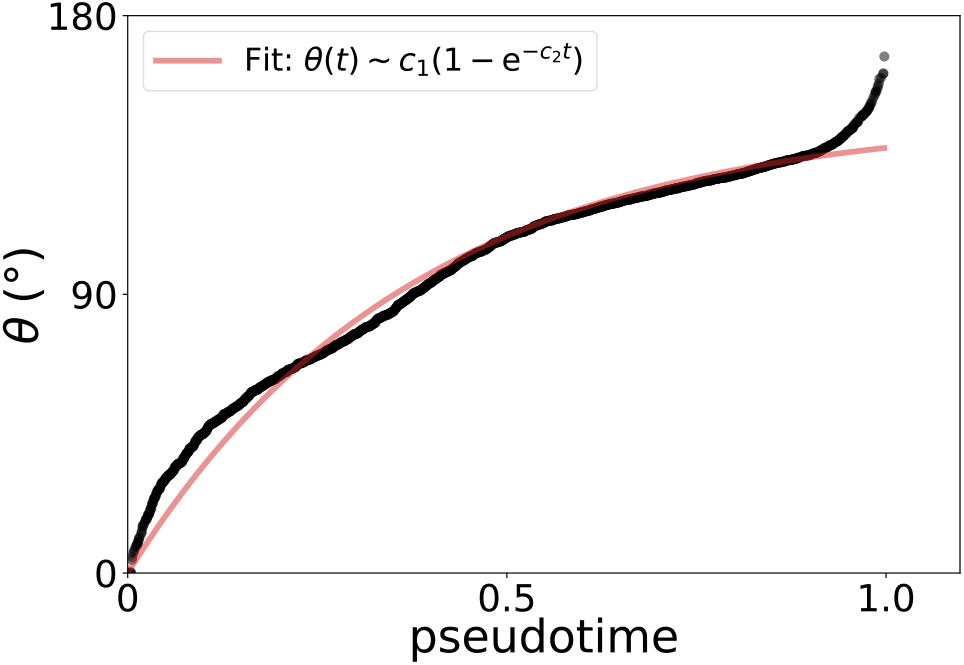
Invagination angle *θ* as a function of the pseudotime *t.* Red line: the fit to the same data as in Fig. 4B according to Eq. (16) for the linear curvature model.

For small *t*, Eq. (16) can be expanded to *θ*(*t*) ~ (*k*_on_/*R*_0_)*t*. The linear increase of *θ* with *t* does not fit the data for small *t* (cf. Supplementary Figure 9).

